# Synergy and antagonism in a genome-scale model of metabolic hijacking by bacteriophages

**DOI:** 10.1101/2024.12.11.628001

**Authors:** Jordan C. Rozum, William Sineath, Pavlo Bohutskyi, Jordan Quenneville, Doo Nam Kim, Connah Johnson, Angad P. Mehta, James Evans, David Pollock, Wei-Jun Qian, Margaret S. Cheung, Ruonan Wu, Song Feng

**Affiliations:** Biological Sciences Division, Pacific Northwest National Laboratory, Richland, WA, USA; Department of Biological Engineering, Utah State University, Logan, UT, USA; Department of Chemistry, University of Illinois at Urbana-Champaign, Urbana, IL, USA; Advanced Computing, Mathematics, and Data Division, Pacific Northwest National Laboratory, Richland, WA, USA; Carl R. Woese Institute for Genomic Biology, University of Illinois at Urbana-Champaign, Urbana, IL, USA; Department of Biochemistry, University of Illinois at Urbana-Champaign, Urbana, IL, USA; Department of Bioengineering, University of Illinois at Urbana-Champaign, Urbana, IL, USA; Environmental Molecular Sciences Division, Pacific Northwest National Laboratory, Richland, WA, USA; School of Biological Sciences, Washington State University, Pullman, WA, USA; Department of Biochemistry and Molecular Genetics, University of Colorado School of Medicine, Aurora, CO, USA; Department of Physics, University of Washington, Seattle, WA, USA

## Abstract

Bacteriophage auxiliary metabolic genes (AMGs) alter host metabolism by hijacking reactions. Previous studies used functional annotations to infer AMG impacts but neglected propagation effects on global metabolism and phage production. We demonstrate the first integration of AMGs and phage assembly into a genome-scale metabolic model, using a general method applied to the infection of *Prochloroccocus marinus MED4* by P-HM2. We experimentally validate our approach to predicting AMG impact on growth using *cp12* mutations in *Synechococcus elongatus*. We predict that 17 directly hijacked reactions substantially impact over 30% of the metabolism, including carbon fixation, photosynthesis, and nucleotide synthesis. We find that indirect impacts are synergistically and antagonistically coupled and are either phage-aligned—shifting feasible reaction velocities in accordance with maximal phage production—or phage-antialigned. Pareto optimization reveals that phage-aligned reactions limit host growth, while phage-antialigned reactions do not. We provide systems-level insight into AMG perturbations, highlighting how nontrivial cascading effects shape microbial functions.

---

The diverse genotypes of coevolved hosts and phages make predicting their interactions challenging. Understanding the dynamics of host–phage interactions is crucial for predicting and controlling host response and viral replication because viruses, including bacteriophages, rely heavily on their host organisms for replication. During replication, viruses hijack the host’s cellular machinery to synthesize the structural proteins and genomic materials needed to assemble new viral particles. A key aspect of viral replication is acquiring necessary building blocks, such as amino acids and nucleotides (*1, 2*). Additionally, viral replication has significant energy and redox demands, including molecules like ATP and NAD(P)H (*3*). Shifts in host metabolism are needed to meet energy, redox, and material requirements critical for the successful proliferation of viruses. These changes are not only theoretical predictions but also are supported by experimental evidence (*4–6*). For instance, infected cells often exhibit increased uptake and usage of nutrients to meet the demands of viral replication (*7, 8*). This can significantly disrupt host metabolism, leading to observable changes in not only the infected cell’s physiology (*9*) but also in the chemistry of the host’s surrounding environment (*10, 11*).

Previous research has demonstrated that phages require specific nutrient ratios to maximize their replication efficiency (*12*). Thus, the uptake rate of specific nutrients in the host cells can be a limiting factor in phage replication, particularly when phages infect their hosts in environments that are optimal for the hosts. In such settings, the competition for essential nutrients intensifies. Marine cyanophages provide an illustrative example of this nutrient limitation. Some of these phages carry the *phoH* gene, which increases phosphate uptake. This may be beneficial for the phage because its biomass contains a significantly higher phosphorus ratio than that of the cyanobacterial host biomass (*12*). The conservation of the phage *phoH* suggests that phosphate availability is critical for successful cyanophage replication, highlighting a specific nutrient demand that is less pronounced in their hosts. These findings underscore likely adaptive strategies that phages employ to redirect the host metabolism so that the phage can thrive in nutrient-limited environments. Understanding how this redirection occurs can provide deeper insights into phage–host interactions and the evolutionary pressures that shape them in various ecological contexts.

Interactions between the host and phage are shaped by evolutionary pressures on both the host and virus (*13*). This coevolution results in a dynamic landscape in which ecosystem-scale effects are coupled to molecular interactions and where understanding the global (i.e., organism-level) effects of specific molecular interactions is paramount. The interaction between viral nonstructural proteins (i.e., those that are not structural components of virus particles) and host components is one venue for such coevolution. Nonstructural proteins can modulate or reprogram the host’s innate immune responses (*14*), stress responses (*15*), and metabolic functions (*16*). In bacteriophages, genes that encode the nonstructural proteins thought to hijack metabolic reactions (e.g., by catalysis or sequestration) are called *auxiliary metabolic genes* (AMGs).

AMGs are widely studied, especially in cyanophage–cyanobacteria systems. Understanding the metabolic hijacking process in these systems has significant biotechnology implications, including the development of novel strategies for controlling harmful cyanobacterial blooms (*17*), developing phage therapy against antibiotic-resistant bacteria (*18*), and industrial bioproduction (*19*). Recent work has shown that bioproduction by *S. elongatus* PCC 7942 is improved by altering its metabolism through the expression of cyanophage-derived sigma factors, demonstrating the potential of phage-inspired approaches for metabolic engineering (*19*).

Marine viral AMGs are generally classified into two groups: Class I AMGs encode enzymes involved in core cellular metabolic processes, including photosynthesis and carbon and amino acid metabolism, while Class II AMGs encode proteins with peripheral functionalities, including transport, iron-cluster assembly, cofactor synthesis, and stress response (e.g., chaperones) (*20–22*).

To ensure sufficient energy for replication, phage-encoded AMGs modulate host energy production by remodeling photosynthetic light-harvesting and electron flow components through Class I AMGs such as the genes *psbA* and *psbD*, which encode the core photosystem II reaction center proteins D1 and D2 (*23*). Additional genes contributing to optimized photosynthetic energy production include *cpeT*, *ho1*, and *pebS*, which are involved in the synthesis of phycobilisome light-harvesting complexes (*24*). High light-inducible proteins (HLIPs) protect photosynthetic complexes and membranes from photodamage by dispersing excessive light energy (*23*). Furthermore, the phage-encoded Class I AMGs *cp12* and *talC* redirect energy and carbon metabolism toward dNTP production by suppressing CO_2_ fixation (Calvin cycle) and activating the pentose phosphate pathway (PPP) (*25*). The adjustment of nutrient acquisition and metabolism is essential to ensure sufficient nitrogen and phosphorus for phage replication in nutrient-scarce environments. These adaptations involve Class II AMGs including the phosphate starvation-inducible gene *phoH*, as well as several AMGs that rebalance nitrogen compounds toward nucleotide biosynthesis, such as ribonucleoside-diphosphate reductase (*nrdA*), nucleotide pyrophosphohydrolase (*mazG*), and cobalamin synthase (*cobS*) (*26–28*). While these processes involving AMGs are well understood based on their functional annotations and assignment to known metabolic reactions, there is no clear theoretical framework for exploring the effect of hijacking individual reactions or for how these effects propagate and couple to alter the host metabolic system.

Genome-scale metabolic models (*29, 30*) are developed to investigate the metabolism-wide effects of perturbations to genes and reaction rates. Flux balance analysis (FBA) and related techniques provide a parameter-robust way to quantify the impact on the quasi-steady-state distributions of reaction velocities (fluxes) that are optimal for biomass production. Flux balance methods are especially useful in determining which reactions are essential for cell growth (*31*) and for assessing how the metabolic flux redistributes in response to gene knockout-induced reaction stoppage (*29,32*). Incorporating the effects of exogenous gene expression, however, remains a challenge. Some progress has been made by incorporating enzymatic constraints (*33*); however, the necessary parameters can be difficult to measure or estimate, and even state-of-the-art machine learning methods require additional post-inference model-specific adjustment (*34*). To our knowledge, no widely accepted kinetics-free method exists to systematically probe the impact of the exogenous expression of metabolic genes, such as that occurring during phage infection. Additionally, using existing genome-scale metabolic models (*35*) to study viral infection requires specification of the virus’ metabolic requirements (*36*). No genome-scale metabolic model of host–bacteriophage interaction currently integrates both the exogenous expression of metabolic genes and phage metabolic demands.

To address this knowledge gap, we developed a strategy to explore how cyanophages hijack *Prochloroccocus marinus* metabolism by expressing known AMGs as a proof of concept that is transferable to other nonmodel organisms. In this study, our focus is on how cyanophage AMGs affect the quasi-steady-state distribution of metabolic reaction fluxes in a host, both individually and combinatorially. We used a previously published iSO595v7 flux balance model (*37*), which is a refinement of an earlier model (*38*). We curated a list of AMGs identified in the cyanophage P-HM2 (*39*) and extracted their predicted or measured effects on specific reactions. The identified reactions are catalyzed by enzymes that are either encoded by AMGs or sequestered by AMG-encoded proteins. We call such reactions *AMG-hijacked* and aim to understand their global impact on the host’s quasi-steady-state metabolic flux distribution and viral replication using FBA and related techniques. To achieve this, we constructed a phage biomass function from the P-HM2 genome and the stoichiometry of its structural proteins by adapting the procedure of (*36*). We integrated the P-HM2 biomass function into the iSO595v7 model of (*37*). Using this enhanced model, we examined which AMG-induced metabolic alterations may favor phage replication over host growth and identified the unique combinatorial effects of AMG-hijacked reaction modulation. These results provide new insight into how phages reprogram host metabolism to boost replication, highlighting mechanisms that drive host–phage interactions and shape their coevolution.

## Results

### Integration of AMGs and P-HM2 phage biomass formation into a genomescale model

From the literature (Table 1), we curated 14 AMGs present in P-HM2 a T4 dsDNA cyanophage that infects *Prochloroccocus marinus* (*40*), and extracted the 17 reactions within the iSO595v7 model that they hijack using annotations from the KEGG database (*41*). The reaction expressions and the subsystems to which they belong are listed in Supplementary Table S1, and additional background information is provided in Supplementary Text. We also constructed a P-HM2 biomass reaction to capture the metabolic demands of phage replication by accounting for the nucleotide synthesis required to replicate the phage genome and the stoichiometric ratios of amino acids in phage structural proteins (see Methods). We calculated that the mass ratio of DNA to structural proteins in the phage particles for the P-HM2 phage is 0.66 (0.399 g DNA/0.6 g protein) and that the the mass ratio of mRNA to structural proteins is 0.022 (we assume that the mRNA is not packed in the phage heads and is recycled by the host metabolism) (Supplementary Table S3). When unconstrained by nutrient abundance, the maximum phage growth in this model occurs at a rate of 0.151 grams per gram of host dry weight per hour. The maximum host growth rate in this model under the same conditions is an exponential growth rate of approximately 0.098 grams per gram of host dry weight per hour (in agreement with the model of (*38*) on which it is based). We compare the demands of the phage biomass to those of the host biomass in Supplementary Table S3. We determined which genes and reactions are required for host or phage biomass production in our model in the standard way using flux variability analysis (FVA; see Methods) (*42*): nonessential reactions are those that contain a flux value of zero within their FVA range when the biomass is constrained to be greater than 1% of its maximum value. We found that 372 of 595 genes (376 of 994 reactions) are essential for host growth^1^. In contrast, only 172 metabolic genes (101 reactions plus the phage biomass reaction) are essential for phage biomass production in this model. All phage-essential genes and reactions that we identified are also essential for host growth (with the exception of the phage biomass reaction). These smaller numbers of genes and reactions that are essential to the phage compared to the host are partly due to the lack of lipids in P-HM2 phage particles, meaning that much of the host’s essential lipid metabolism is not needed to assemble the P-HM2 phage particles. Of the 17 curated AMG-hijacked reactions, nine (PSIIabs^2^, FAKEOrthophosphateEX^3^, R01523, R02017, R02019, R02024, R05223, R05817, and R05818) are essential for host biomass production, and six (PSIIabs, FAKEOrthophosphateEX, R02017, R02019, R02024, and R01523) are essential for phage production. We repeated the essentiality analysis with a lower cutoff value of 10^−7^% biomass production and obtained the same results. The essential genes that we identified in our modified model are consistent with those of the baseline iSO595v7 model of (*37*) but represent a larger fraction, approximately 63%, of metabolic genes than the 47% reported by (*38*) for the iJC568 model upon which iSO595v7 is based. The differences between these models stem from the decoupling of biomass formation and glycogen storage, the refinement of selected reactions, improved gap filling, and additional transport reactions (*37*).

**Table 1:** P-HM2 AMGs identified in the literature. The AMG’s name is given in the first column. The affected reaction ID is given in the Reactions column. In two cases (FAKEOrthophophateEX and PSIIabs), the genes impact metareactions in the model (related to phosphate exchange and photosystem II, respectively)—all other reactions are indicated by their KEGG ID. The reported effect of the AMG on the affected reaction is noted in the Direction column.

| AMG name | Reactions | Direction | Reference | Note |
| --- | --- | --- | --- | --- |
| *cobS* | R05223 | Increase | (43) | Biosynthesis of cobalamin (vitamin B12), important for nucleotide metabolism. |
| *cp12* | R01063, R01523 | Absolute Decrease | (25) | Calvin–Benson cycle inhibitor by sequestering PRK and GAPDH enzymes. |
| *cpeT/cpcT* | R05818 | Increase | (44) | Facilitates fast phycobiliprotein assembly during phage infection. |
| *hlip* | N/A | N/A | (45) | Host survival under moderately high light intensities (not used). |
| *hsp20* | N/A | N/A | (46) | Heat shock response protein (not used). |
| *ho1* | R00311 | Increase | (47) | Phycobilin synthesis. |
| *mazG* | R00426, R00662, R01855, R02100 | Increase | (28) | Proposed to be involved in host genome degradation to reuse building blocks for phage genome replication. |
| *nrdA* | R02017, R02018, R02019, R02024 | R02018 Decrease;<br>All others Increase | (25) | Codes for ribonucleotide reductase to promote dNTP production. |
| *pebS* | R05817 | Increase | (44) | Phage gene *pebS* integrates the functions of *pebA* and *pebB* involved in phycoerythrobilin biosynthesis. |
| *phoH* | FAKEOrthophosphateEX | Decrease | (12) | Promotes phosphate uptake (modeled implicitly in nutrient scenarios). |
| *psbA/psbD* | PSIIabs | Increase | (48) | The genes *psbA* and *psbD* code for D1 and D2 photosystem II proteins. |
| *talC* | R01827 | Increase | (25) | Facilitates enhanced flux through the pentose phosphate pathway for augmented production of NADPH and ribose 5-phosphate. |

We simulated the redistribution of the metabolic flux required to maximize phage particle production to determine the target ranges for hijacked reaction fluxes. We conducted FVA on the iSO595v7 MED4 model (*37, 38*) in a nutrient-rich environment so that RuBisCO efficiency, rather than nutrient uptake, was the growth-limiting factor. We considered two independent scenarios: a preinfection scenario in which the host biomass is optimized, and a post-infection scenario in which the phage biomass is optimized. For each AMG-hijacked reaction, we compared the ranges that allow at least 50% optimal host or phage biomass production (50% is the highest fraction of maximum growth simultaneously achievable by the host and phage in a nutrient-rich environment). We identified one of the endpoints of each range, or zero, to be a target point for each hijacked reaction in accordance with the literature (Table 1). To allow flexibility in combinatorially hijacking reactions, we introduced a small tolerance of 10% of the FVA range in the target flux value. In subsequent results, we constrained individual AMG-hijacked reactions to fall within these target ranges (depicted visually in Figure 1). A majority (13/17) of the AMG-hijacked reaction FVA ranges are largely unaffected (greater than 50% overlap of intervals) by the choice to optimize for the host or phage biomass. For example, R01063 (hijacked by phage *cp12*) has a minimal impact on the host or phage biomass, and its feasible range covers the entire region of the FVA search (±1000 mmol gDW^−1^ h^−1^). Exceptions include two reactions that are hijacked by *nrdA* and involved in nucleotide metabolism. The responses of two reactions, R05818 (hijacked by *cpeT/cpcT*) and R05817 (hijacked by *pebS*), are counterintuitive. These reactions are both thought to have increased flux when their hijacking genes are expressed (*44, 48*); however, the optimal flux through the phage biomass function instead requires a lower flux through these two reactions. In these two instances, we have selected the high-flux portion of the phage-optimal range as the target range.

**Figure 1:**
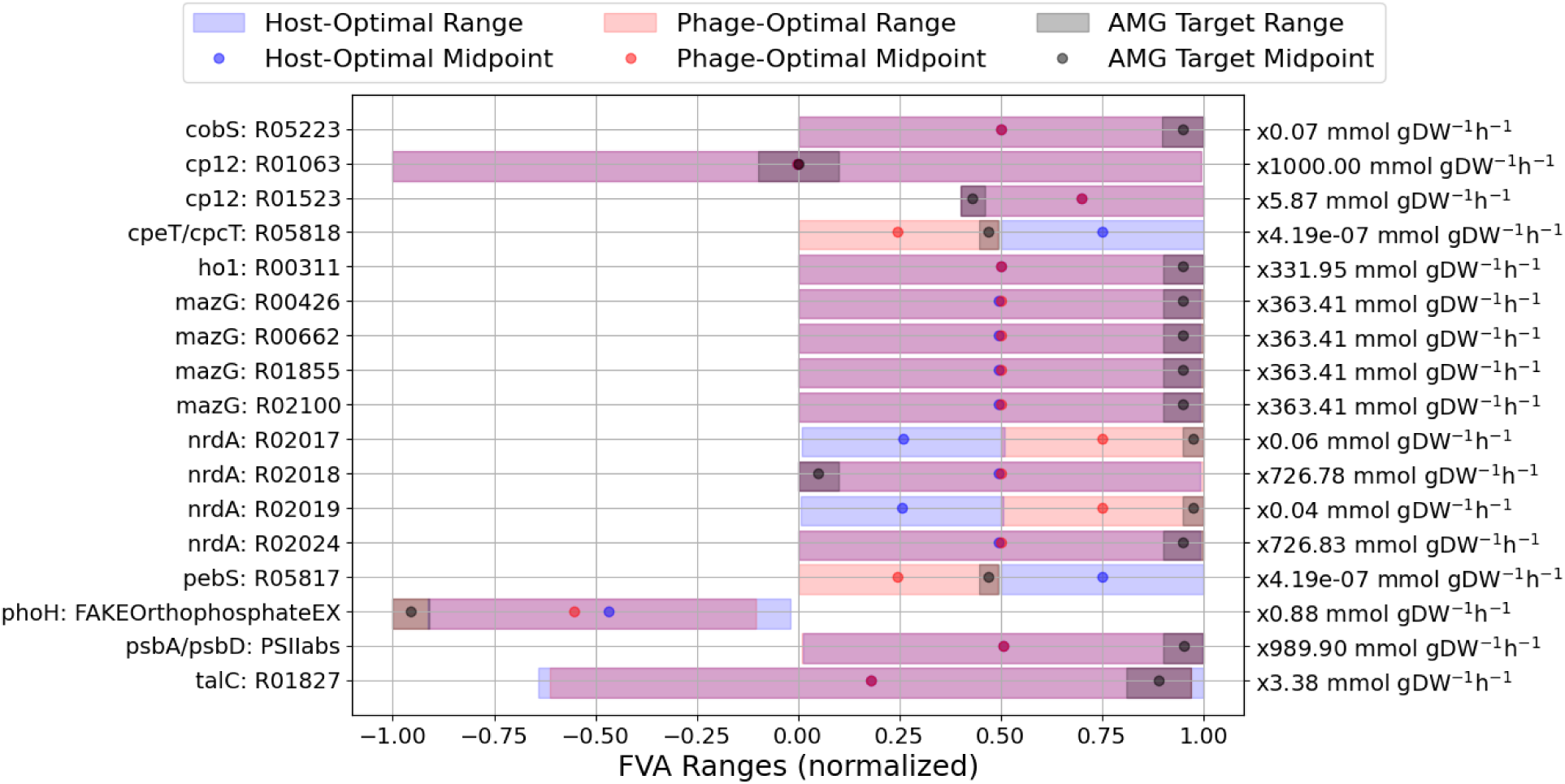
FVA ranges for AMG-hijacked reactions. Blue ranges indicate the flux variability when maximizing the host biomass, while red ranges are for the variability associated with maximizing the phage biomass; any overlap is shown in purple. A decrease in the optimal biomass production of up to 50% is permitted. Ranges are normalized by the most extreme feasible flux across the two scenarios (normalization factors depicted along the right-hand side of the figure). Reaction IDs and subsystems are given along the left-hand side. Using the predicted direction of each reaction flux change as reported in the literature (Table 1), we select 10% of the phage-optimal FVA range to designate as an AMG target range (black).

### The global impact of hijacking a reaction either aligns or antialigns with phage production

We compared the FVA-informed impact of maximizing the host or phage biomass on the global host metabolism. We compared the FVA ranges for each reaction by computing the shift in the FVA midpoint. The largest 30 FVA range shifts, normalized by the largest feasible flux magnitude for each reaction, are shown in Supplementary Figure S1. Affected reactions belong to many metabolic sub-systems including transport and exchange, pyruvate synthesis, glycolysis/gluconeogenesis, carbon fixation, the citrate (TCA) cycle, and various amino acid, nucleotide, and lipid metabolic pathways, consistent with the prior literature on the wide-ranging metabolic impacts of P-HM2 infection in MED4 (*25, 49*). We found that ammonia and orthophosphate uptakes (AmmoniaTRANS, AmmoniaEX, OrthophosphateTRANS, and FAKEOrthophosphateEX reactions) are among the most sensitive to changes in host/phage biomass optimization: optimal phage production required approximately double the rate of ammonia uptake and six times the rate of phosphate uptake required for optimal host growth.

To assess the interplay between phage biomass production and our set of curated AMGs, we fixed the flux of each AMG-hijacked reaction in turn to observe its downstream effects on other reactions when maximizing host growth. We set each AMG-hijacked reaction (one at a time) to the midpoint of its target range (see Figure 1). The FVA midpoints of 321 out of 995 reactions were shifted by more than 0.25 mmol gDW^−1^ h^−1^ in at least one instance because of the fixed AMG-hijacked reaction flux. We compared the host-optimal FVA shifts due to AMG hijacking with the shifts induced by optimizing the phage biomass in the absence of AMGs (Figure 2). Principal component analysis (PCA) identified three distinct categories of AMG-hijacked reactions, based on the loadings of the first principal component, 𝑃𝐶0, which explains 78.7% of the variance among the 17 phage scenarios and the maximum phage biomass scenario. We have named these categories according to how similar their downstream impacts are to the maximum phage biomass scenario (𝑃𝐶0 = 0.26): phage-aligned (𝑃𝐶0 > 0.1; R05223, R01523, R05818, R02017, R02019, R05817, FAKEOrthophasphateEX, and R01827), phage-antialigned (𝑃𝐶0 < −0.1; R00311, R00426, R00662, R01855, R02100, R012018, R02024, and PSIIabs), and minimal impact (−0.1 ≤ 𝑃𝐶0 ≤ 0.1; R01063).

**Figure 2:**
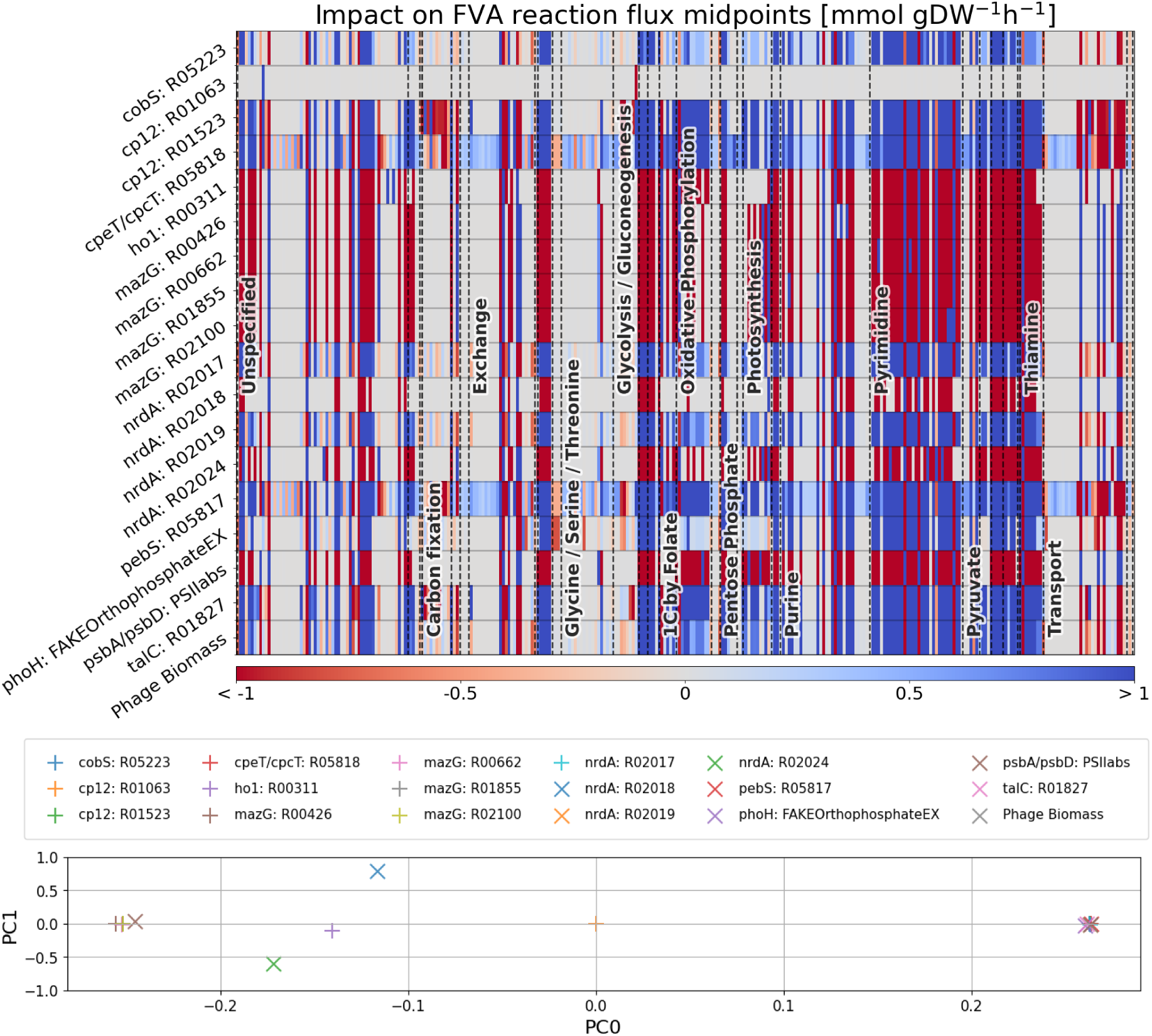
Indirect AMG impact on reactions. (Top) The AMG-hijacked reaction in each row is set to the midpoint of its target range (see Figure 1); in the last row, the phage biomass is instead constrained to its maximum value. Except in this last row, the host biomass is kept fixed for a more direct comparison to healthy cells. The impact on each system reaction (columns) is measured from the midpoint of the FVA range and shown in color (red for decreased flux, blue for increased flux). Affected reactions (columns) are only shown if affected by more than 0.25 mmol gDW^−1^ h^−1^ in at least one instance. They are arranged by metabolic subsystem, and subsystems with more than five affected reactions are labeled. (Bottom) PCA loadings of the AMG impacts in the top panel (including impacts on reactions that are not shown because they fall below the 0.25 mmol gDW^−1^ h^−1^ cutoff).

Hijacking a phage-aligned reaction tends to increase the flux through other reactions in the model when maximizing the host biomass. In contrast, hijacking phage-antialigned reactions tends to have the opposite effect. Though this global trend holds in most subsystems of the model, we found that carbon fixation is an exception: hijacking phage-aligned reactions tends to decrease (rather than increase) the flux through carbon fixation reactions, which are largely unaffected by phage-antialigned hijacking. Consistent with previous work (*25, 49*), we found that *cp12* hijacking of R01523 redirects the metabolic flux from carbon fixation to the PPP and nucleotide synthesis (purine and pyrimidine metabolism) and also that *talC* hijacking of R01827 promotes the metabolic flux through the PPP.

### The trade-off between the host and phage biomasses is modulated by phage-aligned AMGs

In our model, phage-antialigned reactions tend to shift the range of impacted reaction fluxes to lower (or more negative) values, in contrast to phage production, which favors flux ranges with higher midpoints. However, there is no mathematical requirement that the resulting optimal ranges not overlap, in which case maximum phage growth is still possible. Similarly, there is no mathematical requirement that phage-aligned reactions not inhibit phage replication. To assess whether these mathematical possibilities are realized, we examined the impacts of hijacking phage-aligned and phage-antialigned reactions on the host–phage biomass trade-off. First, we constructed the baseline Pareto front for the multiobjective maximization of the phage and host biomasses under various nutrient stresses (e.g., N and P availability) and without hijacking any reactions (left panel of Figure 3).

**Figure 3:**
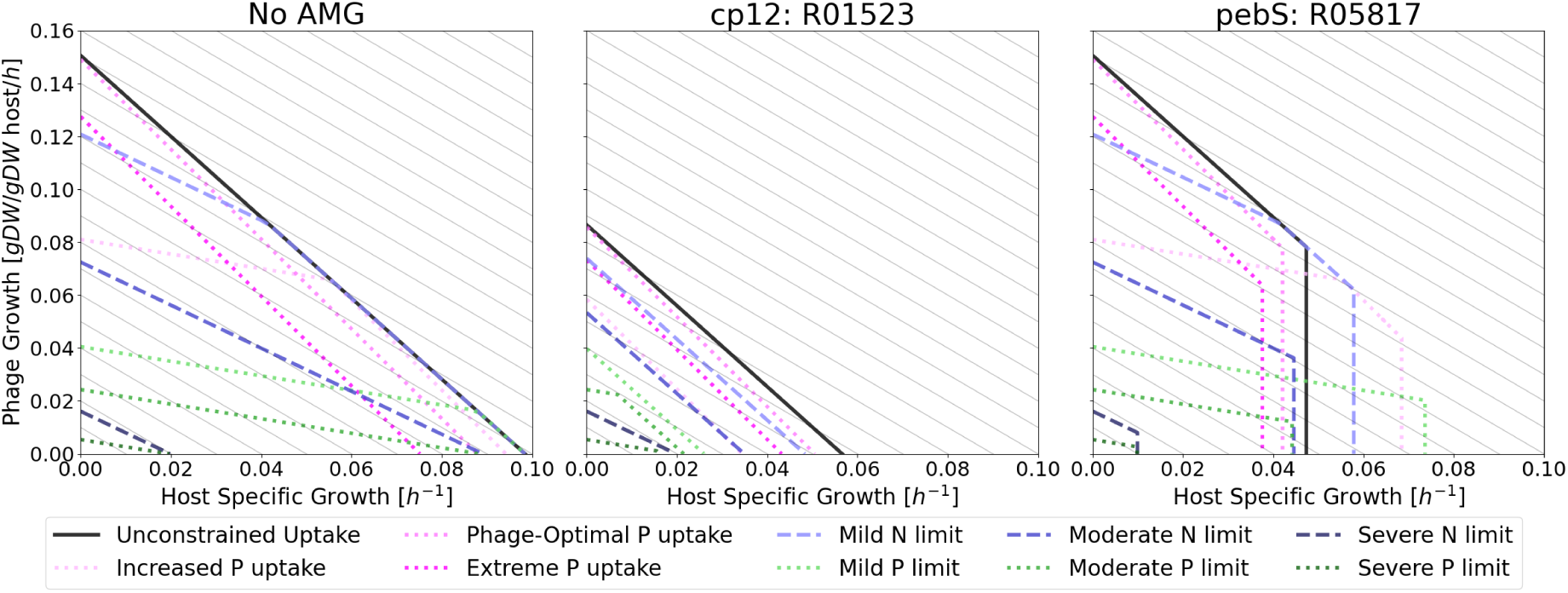
Simultaneous maximization of phage production and host biomass flux. Each panel depicts the instantaneous relationships between the maximum (exponential) host specific growth rate and the maximum (linear) phage growth rate for either no AMG-induced hijacking (left) or for two AMG-hijacked reactions (center, right), as indicated in the figure titles. We show Pareto fronts for several nutrient-limiting scenarios: unconstrained uptake and mild/moderate/severe limiting of N or P uptake. Mild, moderate, and severe limits correspond to 150%, 90%, and 20% of the values required for maximal host biomass production, respectively. When growth is not constrained by nutrient uptake, it is instead limited by the RuBisCO efficiency. In addition, because the gene *phoH* is thought to promote increased P uptake (modeled here as hijacking the FAKEOrthophosphateEX exchange reaction), we considered two scenarios in which the cell is forced to uptake more phosphate than is optimal (Increased P uptake: 3× host-optimal uptake; Phage-optimal P uptake: 6× host-optimal uptake; Extreme P uptake: 12× host-optimal uptake). We provide the remaining 14 AMG-impacted trade-off curves in Supplementary Figure S3.

Based on the insights from previous studies (*12, 37, 38*) and the large increase in N and P uptake rates that we observed when maximizing the phage biomass (Figure S1), we selected N and P availability as the two nutrient stressors. Consistent with these results, both phage and host growth rates are compromised in the model when N or P is limited (compare solid and dashed or dotted lines in the leftmost panel of Figure 3). However, maximum phage growth is more severely impacted in the model: in the moderate N limit scenario (90% ammonia uptake compared to host-optimal), maximum phage growth is 52% lower, while maximum host growth is only 9% lower. Similarly, in the moderate P limit scenario (90% phosphate uptake compared to host-optimal), maximum phage growth is 84% lower, while maximum host growth is 10% lower. Additionally, in all nutrient stress scenarios, the slope of the Pareto front is less negative than that in the unconstrained uptake scenario, meaning that an increase in the host growth rate has a smaller impact on phage production under nutrient stress in the model.

Motivated by prior observations of increased P uptake during infection by cyanophages (*12*), we considered the effects of P uptake rates above the host-optimal level. Fixing the P uptake at 300% of the host-optimal level (increased P uptake scenario) results in a slightly decreased optimal host growth rate (4%). In this scenario, maximum phage growth is reduced by 46% because this increased P uptake rate is only approximately half of the phage-optimal P uptake rate. At a phage-optimal P uptake rate, host growth is 11% lower. Beyond these levels (double the phage-optimal rate), the maximum phage and host growth rates are 15% and 24% lower, respectively. Thus, the host and phage have nonoverlapping ranges of optimal P uptake rates, but both have a wide range (spanning more than an order of magnitude in the case of the host) of suboptimal P uptake rates that allow maximum growth rates above 50% of the unconstrained maximum growth rate.

To study how AMGs might affect the trade-off between host growth and phage production, we constrained each AMG-hijacked reaction to lie within a target range (recomputed as in Figure 1 under each nutrient scenario) and reconstructed the host–phage biomass trade-off curves. All phage-antialigned reactions (Figure 2) preserve the Pareto front. Despite the poor alignment between their indirect impacts on reaction FVA ranges and the requirements of phage biomass production, they do not prevent optimal phage production in any scenario.

All phage-aligned reactions substantially alter the phage–host biomass trade-off in the model. Reactions R05223 (*cobS*), R01523 (*cp12*), and R01827 (*talC*) shift the Pareto front inwards toward the origin, reducing the maximum growth rate for both the phage and host. Additionally, R01523 and R01827 make the Pareto front steeper in some nutrient-limiting scenarios, favoring phage growth (middle panel of Figure 3). The remaining phage-aligned reactions leave the phage-favored upper-left part of the Pareto front unchanged, while rendering the host-favored lower-right portion inaccessible (right panel of Figure 3). In these cases of phage-aligned AMG-hijacked reactions, *optimal host growth is no longer possible, whereas optimal phage growth remains possible*.

### AMGs have both synergistic and antagonistic interactions

Because altering individual AMG-hijacked reactions can shift the midpoints of feasible ranges for other reactions in opposite directions, it is important to understand how these competing effects are reconciled within the host metabolism. To investigate this, we fixed the biomass production at no less than half its maximal value and maximized the total flux impact on AMG-hijacked reactions while simultaneously minimizing the total absolute flux through other gene-regulated reactions by incorporating the standard parsimonious FBA objective as a constraint on enzyme usage. By introducing a linear *AMG reaction emphasis factor*, denoted 𝛾, to weight these two flux objectives (see Methods), we simulated the trade-off between prioritizing AMG flux impacts and reducing the overall flux. Higher values of 𝛾 favor AMG impacts, while lower values prioritize reducing the total absolute flux (as in parsimonious FBA). We show the resulting change in reaction fluxes, compared to a baseline parsimonious FBA distribution, for the 20 most highly affected reactions in Figure 4, alongside the change in AMG-hijacked reaction fluxes that were obtained during this procedure.

**Figure 4:**
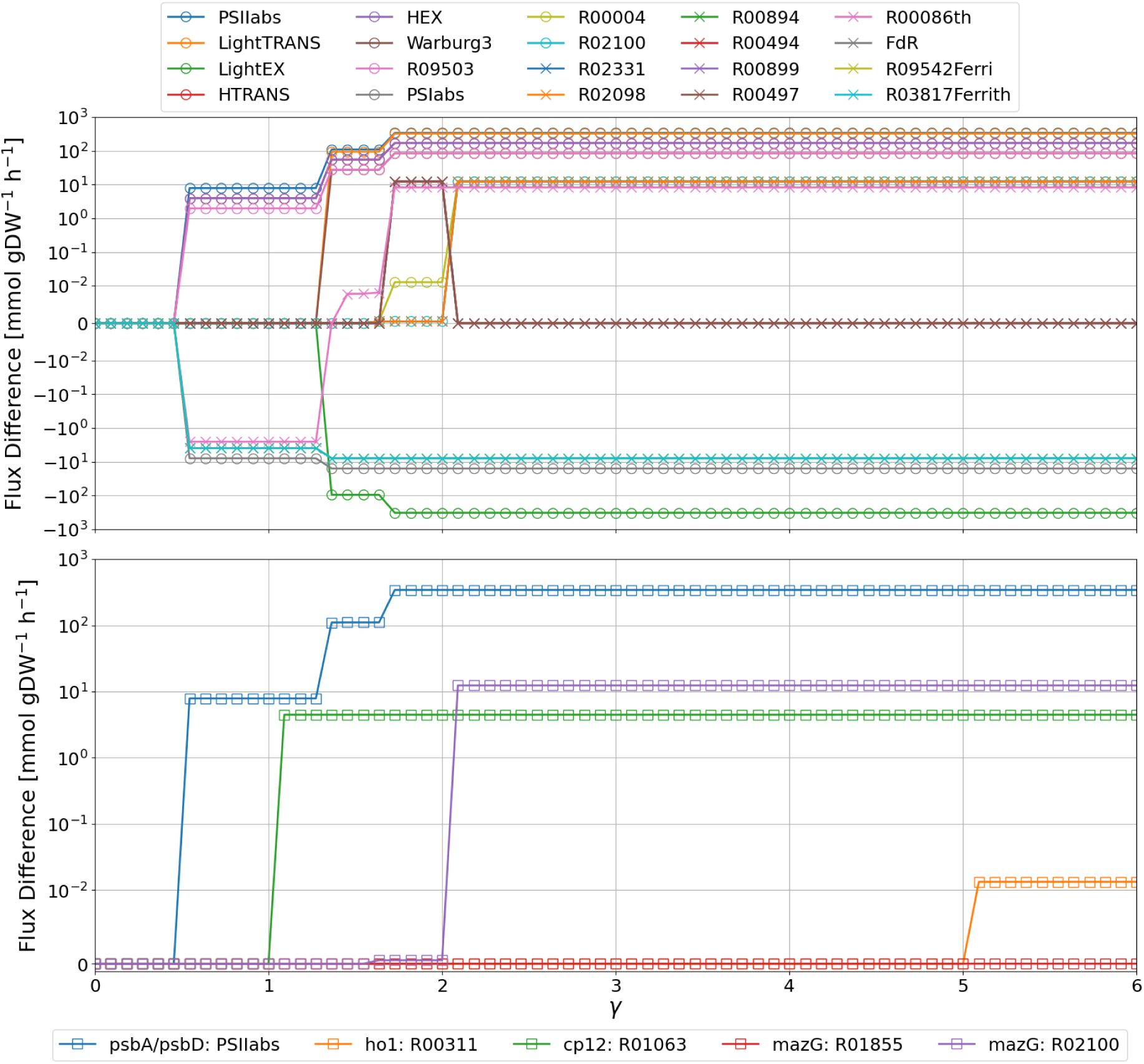
Changes in reaction fluxes relative to a parsimonious FBA baseline. Flux change is shown as a function of AMG reaction emphasis factor, 𝛾. Increasing 𝛾 more heavily weights changing the flux of AMG-hijacked reactions relative to minimizing the flux through other gene-regulated reactions (see Methods). In the top panel, the 20 reactions with the largest responses are shown. The ten with the largest responses are marked with circles; the remaining ten use an X symbol. In the bottom panel, AMG-hijacked reactions with nonzero flux change are shown. The vertical axes use a symlog scale, so that the region between ±10^−2^ mmol gDW^−1^ h^−1^ is linearly scaled, while the regions outside this range use a logarithmic scale.

We also considered the impacts on specific mass fluxes between metabolic subsystems (see Methods). We analyzed the changes in the specific mass flux through each subsystem using mass flux difference (MFD) graphs for the most affected subsystems. These graphs highlight the change in the mass flux through and between metabolic subsystems and are defined in Methods. MFD graphs for four values of 𝛾 are depicted in Figure 5. In Supplementary Figure S2, we plotted the change in the mass flux through the 20 most affected subsystems as a function of 𝛾.

**Figure 5:**
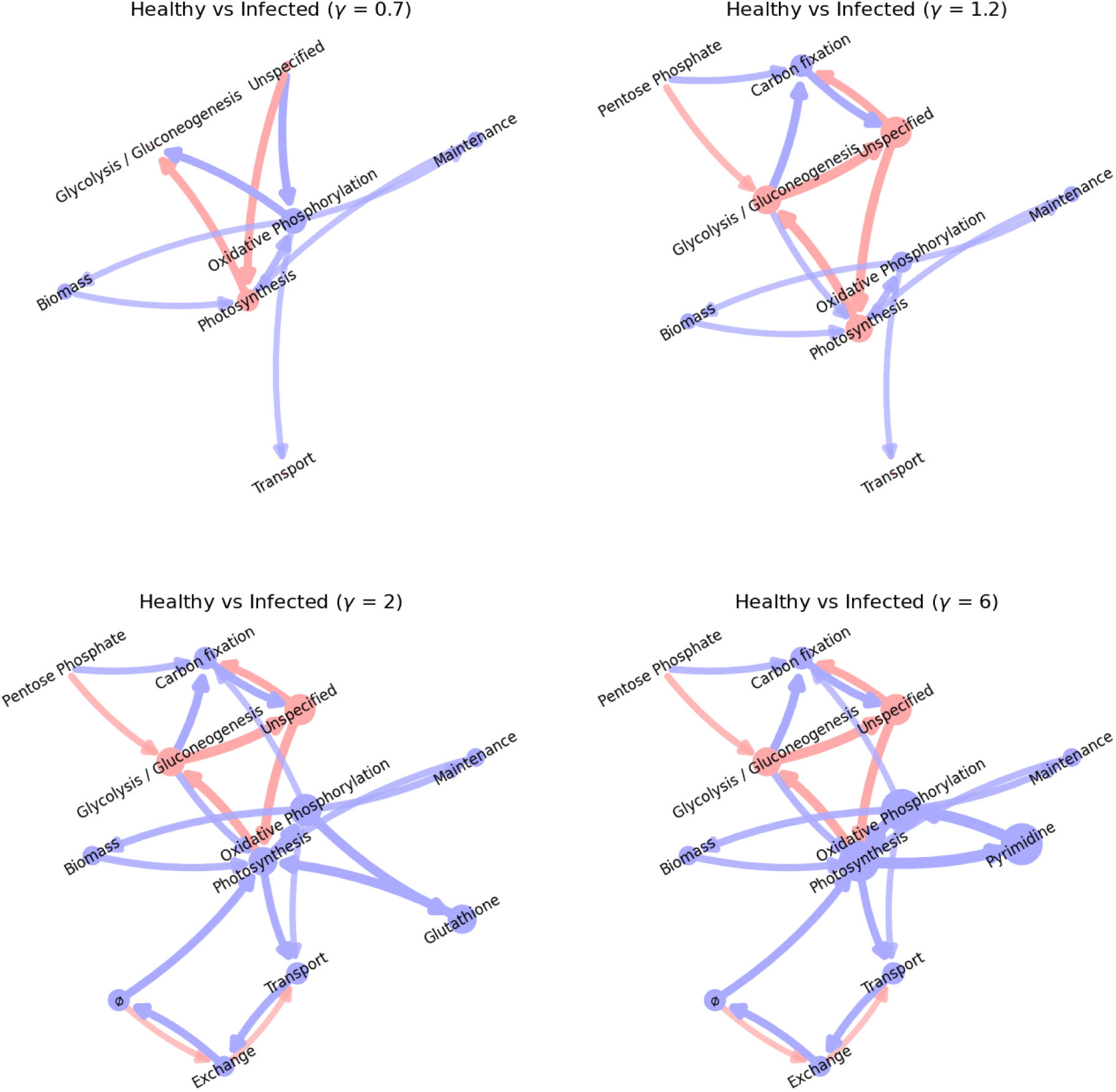
Changes to subsystem-specific mass flux graphs. Flux changes are represented as MFD graphs (see Methods) at various values of the AMG reaction emphasis factor, 𝛾. The node size and edge thickness and opacity are scaled according to the absolute change in the specific mass flux, with blue indicating an increase and red indicating a decrease in the infected system fluxes relative to the uninfected fluxes. Node positions are fixed across the four graphs. Edges with specific mass flux changes below 0.01 h^−1^ are not shown, nor are nodes that lack incoming or outgoing edges above this threshold.

For small values of 𝛾, only the photosynthetic reaction PSIIabs (*psbA*/*psbB*) is directly affected (bottom panel of Figure 4). In our numerical experiments, increasing the flux through this reaction decreased the mass flux through photosynthetic pathways and increased the mass flux through the oxidative phosphorylation subsystem (top left panel of Figure 5). For 𝛾 > 1, the suppression of R01063 by *cp12* (increased negative flux through this reversible reaction) becomes favorable (top right panel of Figure 5). Setting 𝛾 = 2 permits R01855 hijacking by *mazG* and a further increase in the PSIIabs flux (bottom left panel of Figure 5). For this value of 𝛾, the mass flux through both the photosynthetic and oxidative phosphorylation pathways increased relative to the baseline, with more flux diverted to the biomass and carbon fixation subsystems and away from glycolysis and gluconeogenesis. Beyond 𝛾 = 2, R01855 was hijacked by *mazG*, leading to a large (1.95 h^−1^) increase in the specific mass flux through purine metabolism (bottom right panel of Figure 5).

For 𝛾 ≥ 5, R00311 was hijacked by *ho1* in our simulations, but this had a negligible impact on the subsystem mass flux (Supplementary Figure S2). All of the increased reactions belong to the phage-antialigned group identified in Figure 2. This may be due to the tendency of this group of reactions to decrease the flux globally, which is favored when minimizing enzyme usage.

We further explored the synergistic and antagonistic effects of AMG-hijacked reactions by fixing pairs of AMG-hijacked reaction fluxes to their target ranges (Figure 1 and Supplementary Figure S4). To assess the interdependence of hijacked reaction pairs, we compared the edge-wise sum of MFD graphs computed from each hijacked reaction individually to the MFD graph obtained when hijacking both reactions simultaneously. We quantified this comparison using the edge-wise root mean square difference (RMSD) between these two graphs (Figure 6). In some cases, the two target ranges for the AMG-hijacked reactions are incompatible, resulting in infeasible solutions. Even reactions hijacked by the same gene can have this property, such as R00426 and R00662, both hijacked by the gene *mazG*.

**Figure 6:**
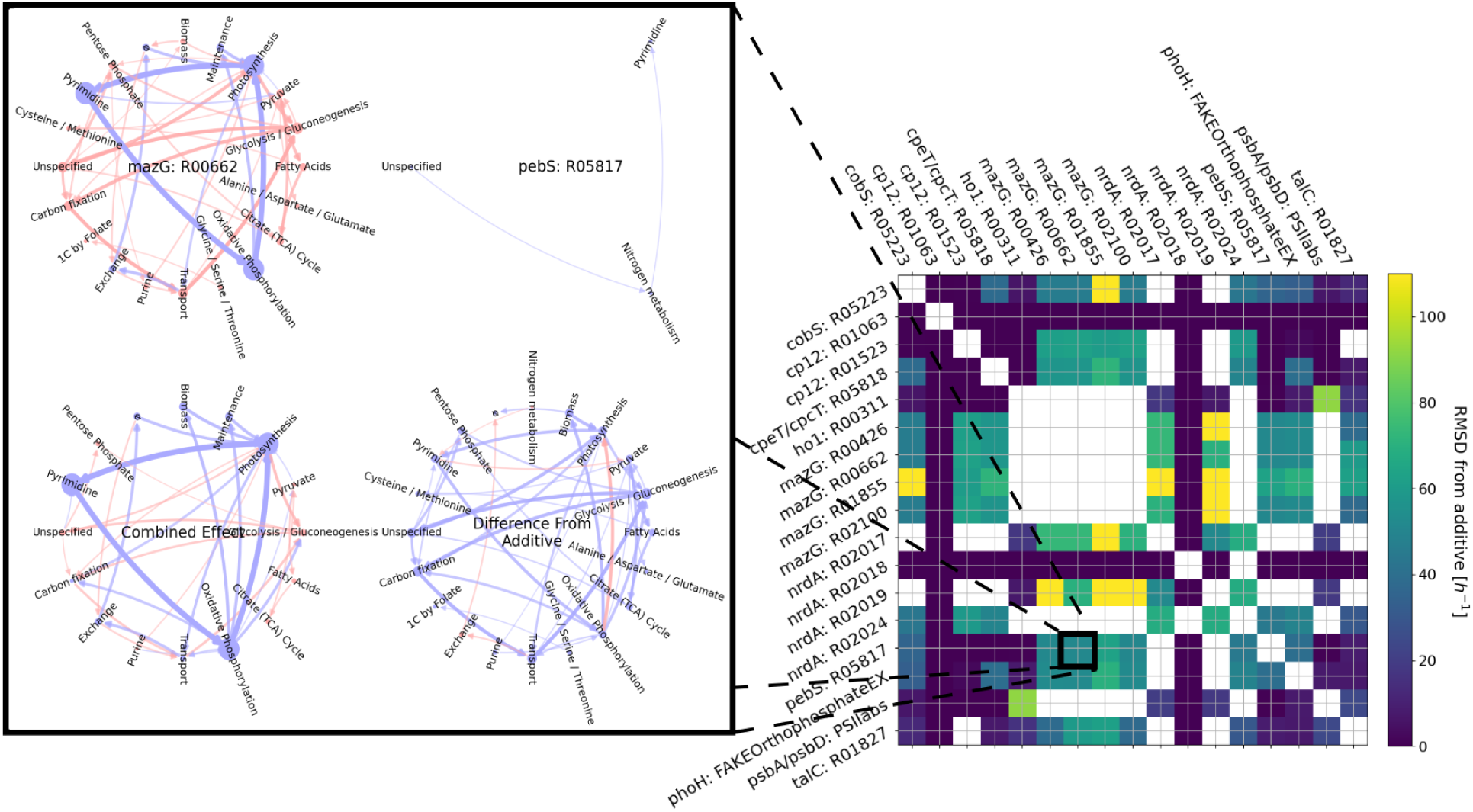
Interdependence of subsystem mass flux response to AMG-hijacked reactions. Each entry of the heat map matrix corresponds to a pair of AMG-hijacked reactions, indicated by the row and column labels. For each pair, two combined MFD graphs were computed: the true MFD graph obtained from constraining both hijacked reactions to their target ranges (see Figure 1) and the additive MFD graph obtained by adding (edge-wise) the MFD graphs obtained by individually hijacking each reaction. The nonadditivity is the edge-wise RMSD of the additive and true MFD graphs. Off-diagonal white cells indicate that the combined hijacking of the two AMGs did not yield a feasible solution. An example with a moderate RMSD comparing the MFD graph for individual and combined effects of hijacking two reactions is shown in the inset. In all graph plots, edges with specific mass flux changes below 0.01 h^−1^ are not shown, nor are nodes that lack incoming or outgoing edges above this threshold.

In some scenarios, we observed antagonistic effects. The inset of Figure 6 highlights how *pebS*, via R05817, moderates the impact of *mazG* via R00662 in the model. At the system level, R05817 has only a small impact on the mass flux on its own: only three subsystems have incoming or outgoing flux changes larger than 0.01 h^−1^. In contrast, R00662 impacts 20 subsystems above this threshold, with the largest impacts on the mass flux through the Pyrimidine, Photosynthesis, and Oxidative Phosphorylation subsystems. The flux is increased in all three subsystems when R00662 is hijacked. The largest inhibitory impact of R00662 hijacking in the model is on Glycolysis/Gluconeogenesis. Because the impact of R00662 is much larger than the impact of R05817, their combined effect would be very similar to that of R00662 alone if they acted independently; however, we instead observed that the inhibitory impacts of R00662 on Glycolysis/Gluconeogenesis and several of its neighboring subsystems are substantially decreased by R05817 hijacking (see inset of Figure 6). In contrast, the promotive impacts in the combined R00662–R05817 hijacking scenario on the Pyrimidine, Photosynthesis, and Oxidative Phosphorylation subsystem mass fluxes remain high.

We also noted a > 0.01 h^−1^ increase in the mass flow from the Oxidative Phosphorylation subsystem to Biomass reactions in the combined R00662–R05817 hijacking scenario that does not appear when hijacking either reaction alone.

The most extreme deviations from purely additive effects occur between reactions hijacked by *nrdA* and *mazG*, both of which modulate nucleotide metabolism (Figure 6). Apart from these, the largest deviation from a purely additive effect occurs between PSIIabs and R00311, visualized in Supplementary Figure S5, alongside a summary of the subsystem fluxes that most substantially deviate from an additive response. In the example of PSIIabs and R00311, the largest deviations from a purely additive effect involve Photosynthesis, Oxidative Phosphorylation, Pyrimidine, and Porphyrin/Chlorophyll metabolism. In particular, in the combined hijacking scenario, the mass flux through the Photosynthesis subsystem is synergistically altered so that it favors diverting the mass flux from Oxidative Phosphorylation to Porphyrin/Chlorophyll and then to Pyrimidine metabolism.

### CP12 impacts on cyanobacteria growth are reduced by N scarcity both *in silico* and *in vivo*

Phage *cp12* hijacking the reaction R01523 inhibits host growth in our simulations when N is available but not when it is scarce (compare the left and center panels of Figure 3). To further explore this inhibition, we first computed the ratio between N-unlimited optimal host growth to N-limited optimal host growth for 21 evenly spaced fractions of the unconstrained N flux. We then repeated this computation with reactions R01523 and R01063 simultaneously hijacked by *cp12*. We observed that the ratio of optimal growth rates becomes insensitive to variations in N availability below a threshold that depends upon the P availability and the strength of carbon fixation suppression by the CP12 protein encoded by *cp12*.

Quantifying and validating this effect *in vivo* poses significant challenges in MED4, which is genetically intractable. However, another model cyanobacterium, *S. elongatus* PCC 7942, is well studied and genetically tractable. Thus, we applied our methods to *S. elongatus* PCC 7942 using the previously published genome-scale metabolic model iJB785 (*50*), which we modified to be RuBisCO-limited in a replete nutrient environment as in (*37, 38*). We first verified that carbon fixation is inhibited by *cp12* in both the MED4 and *S. elongatus* metabolic models by hijacking the shared R01523 and R01063 reactions. We also compared the flux adjustments of all shared reactions relative to the *cp12*-inhibited reaction R01523 in both the MED4 and *S. elongatus* models (Supplementary Text), and the highly correlated sensitivities imply similar function and impacts of *cp12* in both MED4 and *S. elongatus* metabolism (Supplementary Figure S6). We then repeated our simulation method to compute the relationship between N availability, *cp12*, and optimal growth in iJB785. The MED4 and *S. elongatus* models predict the same cell growth penalty due to *cp12* as a function of the total N flux (Figure S7). All these *in silico* analyses support the fact that experimental validations of predictions from the *S. elongatus* model are transferable to MED4.

To validate the predicted effect of phage AMG *cp12* on N-limited *S. elongatus* growth, we engineered a mutant strain constitutively expressing P-HM2’s *cp12* and measured growth curves for the mutant and wild type (WT) in 10 N-limiting scenarios. For validation, we consider high- and low-light scenarios (Figure S8). With four replicates of each culture condition, this results in 160 growth curves in total. We determined the maximum growth rates by fitting logistic functions to all 160 growth curves, discarding replicates with large uncertainties in the estimated growth rates. Then, we calculated the log2 fold-change (log2FC) in maximum growth in the mutant strains relative to the WT and compared it with the log2FC in maximum growth predicted from the FBA analysis (Figure 7).

**Figure 7:**
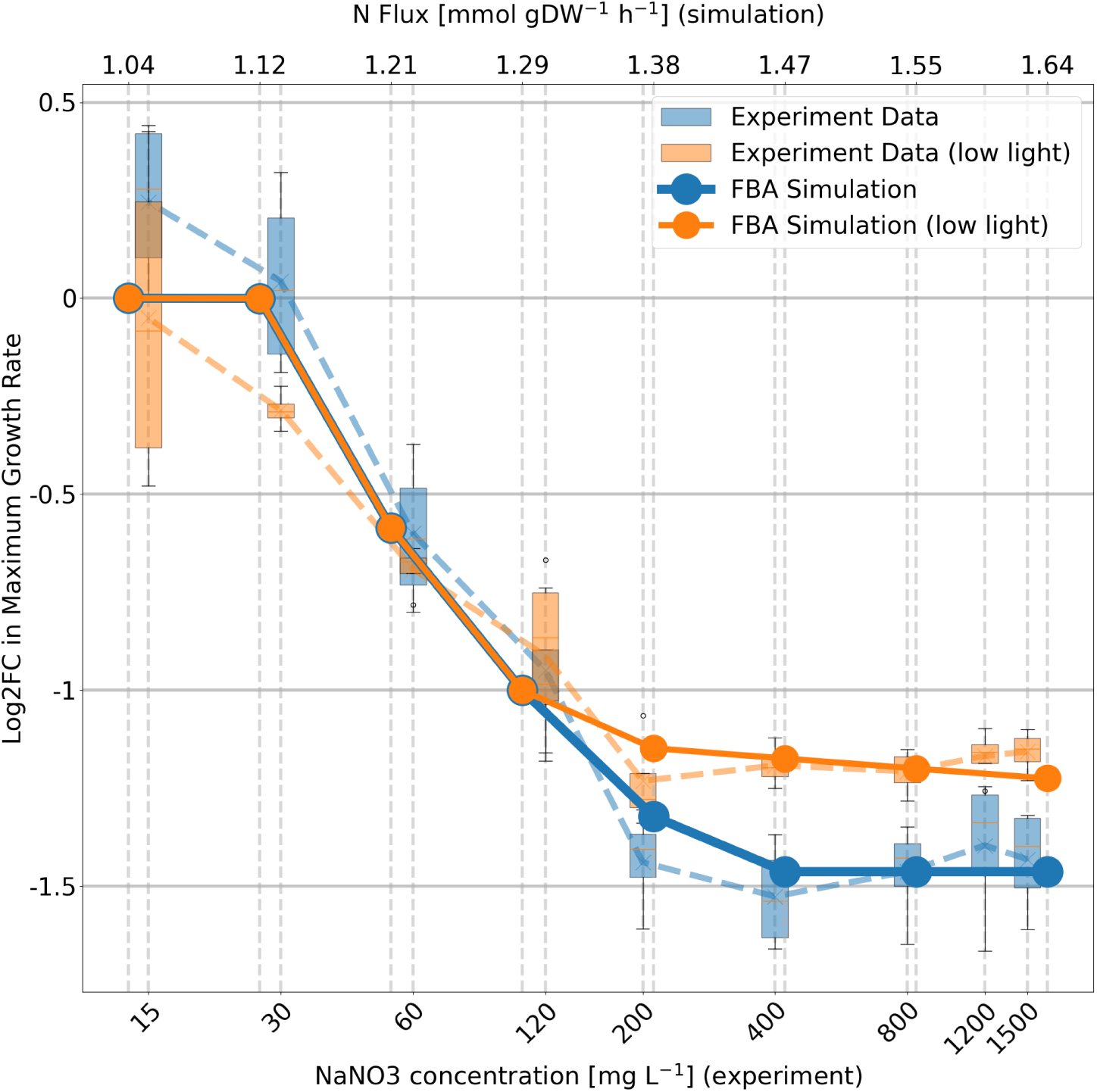
Comparison of the maximum *cp12* mutant and WT growth rates *in silico* and *in vivo*. We depict the log2 ratio of mutant to WT maximum growth on the vertical axis. For the experimental data, we determined growth rates by fitting logistic growth curves to OD_750_ time series with the background subtracted; for the simulated data, growth rates were determined from the biomass flux. We considered high-light (blue) and low-light (orange) conditions and depict nine N-limiting conditions (bottom horizontal axis; note the logarithmic scale in NaNO_3_ concentration). A tenth experimental condition in which no NaNO_3_ was supplied resulted in growth rates indistinguishable from zero and is not depicted here. FBA simulation results are overlaid, with evenly spaced N-flux constraints (top horizontal axis). The low-light condition is simulated as a 40% decrease in the photon flux through photosystem II.

We find good qualitative agreement between simulation and experiment (Figure 7) as well as high correlation (𝑅^2^ = 0.93 for high light conditions and 𝑅^2^ = 0.86 for low light conditions) between the log2FC values of experimental measurements and model simulations (Figure S9). In particular, at low levels of nitrogen availability, both show similar growth rates for mutant and WT strains, while at high N availability, the WT has more than double the maximum growth rate. Both simulation and experiment show a gradual transition between these regimes. We find that the difference in maximum growth rate at high N availability is diminished in low light in both simulation and experiment.

## Discussion

Interactions between bacteriophages and their hosts are complex: they involve the ecosystem-scale exchange of metabolic genes over evolutionary time scales, coupled to minute-by-minute changes in intracellular reaction fluxes (*27, 51, 52*). A critical step in unraveling this complexity and understanding whole-cell response to viral infection is to characterize how exchanged AMGs hijack metabolic reactions to shape host metabolism and promote viral propagation. Genome-scale metabolic models can accelerate these biotechnology applications by predicting optimal combinations of phage-derived AMGs for redirecting metabolic fluxes toward desired products while maintaining cell viability. Such a systems-level understanding of metabolic hijacking mechanisms can inform the rational design of synthetic metabolic circuits that mimic successful viral strategies for metabolic reprogramming. This is a challenging task in part because dynamical parameters are poorly constrained. Fortunately, static methods, such as flux analysis, can uncover important parameter-robust insights. Our approach represents a framework for studying possible AMG impact scenarios using FBA and other closely related techniques. In analyzing the *P. marinus* infection by the cyanophage P-HM2, we uncover the systems-level impacts brought about by the viral hijacking of individual metabolic reactions.

Our approach fills critical gaps in understanding AMG impacts by moving beyond functional annotation, which can overestimate the role of AMGs because of limited validation at the cellular and community levels. Functional annotations of P-HM2 AMGs detected in metagenomes suggest their involvement in key metabolic pathways, such as photosynthesis and carbon fixation (*25, 48*). However, inferring AMG impacts solely from these annotations is insufficient to capture their full biological significance. To address this, we employed a whole-genome-scale metabolic modeling approach, incorporating—for the first time—both AMG perturbations and phage particle production as an objective function. Our phage production objective function predicts that optimal phage biomass production requires substantially more nitrogen and phosphorous uptake than optimal host growth. This aligns with previous observations (*12*) that infected host cells exhibit increased nutrient uptake. Alternatively, this may reflect the phage’s insensitivity to other limiting factors (e.g., the energetic demands of lipid formation), allowing the phage to better leverage additional available nutrients. By probing the impact of AMGs at a genome scale, we identified both phage-aligned and phage-antialigned reactions potentially hijacked by AMGs, revealing metabolic insights that functional annotations alone cannot provide. Additionally, our approach provides new opportunities for predicting and understanding emergent phenotypes driven by AMG-induced resource and energy redistribution when modeling the infected cell as a whole. For example, subsystem flux analyses reveal a redirection of the flux from carbon fixation to the PPP. This is in line with the observation of reduced carbon fixation in marine cyanobacteria upon phage infections (*49*). However, the nonmonotonicity of some of these system-level responses, such as in the photosynthetic flux and oxidative phosphorylation, suggests complex interactions between AMG-driven pathways and host cellular networks. This highlights the importance of systems-level approaches in understanding how AMGs reprogram the host metabolism to optimize resource allocation during infection. By introducing the concepts of phage-aligned and phage-antialigned hijacking, we provide a framework to classify AMG functions and prioritize candidates for experimental testing. Phage-antialigned AMGs, which individually facilitate significant metabolic shifts without altering the host–phage biomass trade-off, may yet significantly impact phage production through their collective interactions and warrant further investigation. To date, there are only a few studies that investigate phage-driven metabolic reprogramming in host cells (*53*) and experimentally validate the enzymic functions of the phage AMGs (*54, 55*). These studies reinforce the need for collective experimental and computational efforts to speed the discovery of the complex roles of AMGs. By integrating insights from these works, we open new avenues for the theoretical exploration of AMGs’ impacts and prioritize AMGs for more in-depth experimental validation.

Though our analysis describes the global effects of combinatorial reaction hijacking, not all AMG-hijacked reactions changed flux in our numerical experiments—in fact, most did not. In some cases, this is because increasing the flux in some of these reactions requires a large increase in the total flux along the pathway of the AMG-hijacked reaction, thus requiring a substantial increase in enzyme production. Even disregarding enzymatic constraints, it is not stoichiometrically feasible to simultaneously maximize the impact of all observed AMGs. Thus, we have considered scenarios in which the individual impacts of AMGs are not maximized. It is possible, for instance, that the produced AMG enzyme is not efficiently used by the host metabolism, and previous work has demonstrated time-dependent transcriptional regulation of AMGs (*27*). Additionally, a host cell may have branch points in its metabolism that are seldom used in tandem. If a phage benefits from hijacking one branch, the cell may divert the metabolic flux toward the alternate branch to compensate. In such cases, the phage may benefit from hijacking reactions along both branches, even when only one branch or the other can carry the flux. In some cases, however, AMGs have antagonistic, competing impacts on metabolic fluxes. We conjecture that these antagonistic interactions, when coupled to the time-varying expression of AMGs in the host organism (such as differentiated by the day/night cycle (*27, 56*)), may play a role in synchronizing lysis for the optimal burst size. This latter explanation is supported by the extent to which hijacking-induced changes to the host-optimal flux ranges either align or antialign with those induced by optimizing the phage biomass.

We validated our approach by comparison with the model organism *S. elongatus* and a mutant *S. elongatus* strain that we engineered to express P-HM2 *cp12*. The *cp12* gene encodes the intrinsically disordered CP12 protein that sequesters key enzymes in the carbon fixation metabolism of photosynthetic organisms. CP12 responds to the cellular redox environment by forming a structure-stabilizing disulfide bond that allows CP12 to bind to the Calvin–Benson cycle enzyme tetramer GAPDH (*57*). PRK, another Calvin–Benson cycle enzyme, can bind to join two GAPDH-CP12 complexes to form the *dark complex* GAPDH_8_CP12_4_PRK_4_ in which the catalytic activities of GAPDH and PRK are severely inhibited (*58*). We replicated our AMG approach in a previously published model of *S. elongatus* and demonstrated that the global effects of *cp12* expression are similar in both models. From our numerical analysis, we predicted that N becomes limiting at a similar value of flux regardless of *cp12* expression. Our simulations indicate that the constitutive expression of *cp12* inhibits growth in a high-N environment but not when N is scarce. Furthermore, in our simulations, the magnitude of this high-N growth inhibition diminishes in low-light conditions. We validated both predictions experimentally. We interpret this effect to indicate that the sequestration of Calvin–Benson cycle enzymes by CP12 inhibits efficient usage of abundant N. Our experimental results not only bolster our confidence in these specific predictions and interpretation, but also in our method for implementing AMG expression in genome-scale metabolic models. Concordant predictions from the *P. marinus* MED4 and *S. elongatus* models regarding CP12’s influence on growth support its evolutionarily conserved function as a regulator protein modulating Calvin–Benson cycle activity and redistributing carbon through alternative pathways. Comprehensive, comparative studies tracking metabolic reprogramming across a wider spectrum of AMGs and varied phage–host systems are needed to clarify the multifaceted roles that these genes play in phage–host coevolution.

Very few of our system-level predictions can be obtained straightforwardly using purely local methods. Instead, we must consider the global metabolic response to AMG perturbations to uncover phage-aligned or phage-antialigned flux changes across multiple metabolic subsystems. Moreover, the cascading effects that give rise to a global response are not necessarily mediated by traditionally recognized core pathways. When plotting metabolic networks, reactions that share only common metabolites, such as ATP, are not connected to avoid excessive visual complexity, but the metabolism is inherently complex. Frequently disregarded connections can become important in characterizing the global response to a phage hijacking nucleotide metabolism, for example, encouraging ATP (and dATP) production at the cost of lipid metabolism or other normally essential processes. Similarly, the competition between subsystems for common metabolites may play an important role in the alteration of the host metabolism; predicting the winners and losers of these competitions requires engaging with the metabolic network as a whole. The complexity only increases when we must untangle the collision of perturbation cascades induced by AMGs hijacking reactions from widely separated portions of the metabolic network. This work has begun to address the complexity of the quasi-stationary response to AMG hijacking, but yet another layer of complexity remains to be incorporated in future work: the temporal heterogeneity of AMG expression (as described by (*27*)). The characterization of the metabolic response we have presented here relies on a computationally efficient and parameter-robust method, laying the groundwork for more resource-intensive in-depth investigations of AMG hijacking response during bacteriophage infection.

## Methods

### Building the P-HM2 phage biomass and AMGs into a genome-scale model

In this study, we used as a starting point the genome-scale iSO595v7 model of *Prochloroccocus marinus* MED4 metabolism, which was released by (*37*) (retrieved from https://github.com/ segrelab/Prochlorococcus_Model) as a refinement of the model developed by (*38*) and which has received several small updates since. In addition to various updates by the original creators of iSO595v7, we have made a small correction to the RNA synthesis reaction so that dNMP (rather than dNTP) polymers are constructed from dNTP, releasing diphosphate in the process. Though in principle this could affect the global phosphate balance in the model, we did not observe any significant change in the system-level behavior as a result of this adjustment. Reactions are annotated with the metabolic subsystem they are assigned to in KEGG (*41*). Metabolites are assigned one of five compartments (extracellular, cytoplasm, carboxysome, thylakoid, or periplasm). This model contains 994 metabolic reactions among 802 metabolites (including duplicates from different compartments). The reaction activity is governed by 595 genes.

To add the phage biomass function to the iSO595v7 model, we identified the stoichiometric requirements for phage growth due to structural proteins and genetic material. We created a dictionary of structural proteins in the P-HM2 phage using previously published protein counts from the homologous and canonical Myoviridae member T4 bacteriophage (*59*). We used protein sequences from the T4 bacteriophage protoeme (proteome ID UP000009087) in a BLAST search against the UniProt P-HM2 proteome (proteome ID UP000006538). We identified each protein with the lowest e-value in the BLAST results as the homologous protein in the P-HM2 virion particle and used the corresponding stoichiometry of T4 structural proteins as the stoichiometry of their homologous structural proteins in P-HM2. All the stoichiometry matching between T4 and P-HM2 and BLAST results can be found in Supplementary Data S1.

Using the stoichiometry dictionary of structural phage proteins as input, we adapted the procedure of (*36*)—designed for computing the biomass of SARS CoV2, a single-stranded RNA virus—to generate the biomass function of P-HM2, a double-stranded DNA virus. First, we used the stoichiometry of structural proteins from the UniProt P-HM2 proteome (proteome ID UP000006538) inferred from the BLAST search results to calculate the number of each type of amino acid required to form a single virion particle. For DNA requirements, we used the P-HM2 genome (ENA accession ENA GU075905.1) to construct the number of each type of nucleotide present in a fully packed virion particle. For mRNA requirements, we assume an average of 540 protein copies per mRNA transcript, in line with previous measurements in bacteria (*60*); in practice, so long as it remains large, the precise value of this ratio has a negligible impact on the maximal phage production rate. We estimated the ATP consumption for amino acid and nucleotide polymerization in each phage using the method of (*36*). From the total molar weight of each component consumed, we computed the molar weight of one P-HM2 phage particle to normalize the biomass expression so that the stoichiometric coefficients describe the number, in millimoles, of each metabolite required to produce one gram of virus. We measure the production rates of these metabolites in millimoles per gram of dry weight per hour. Thus, our model outputs the virus biomass production rate in grams of virus produced per hour per gram dry weight of host. The detailed composition of the biomass function is given in Supplementary Table S2.

This model assumes a full and proper packing of phage heads and that the constituent phage structure proteins are only created in the correct stoichiometric ratios. P-HM2 is known to mispackage host DNA inside its capsid, but the frequency of this occurrence is low and unlikely to significantly impact the model (*61*).

To integrate AMGs into the model, we used previously identified AMGs from (*40*) to conduct a BLAST search against the P-HM2 genome (accession GU075905.1). We then identified the reactions in iSO595v7 that are indicated in the KEGG database (*41*) as most directly impacted by each AMG. To determine the direction (increase or decrease) of the AMG’s impact on the associated reaction, we conducted a literature search of each AMG (the results are summarized in Table 1). Only one AMG that we identified, *cp12*, caused an inhibition of reactions. Its associated protein, CP12, inhibits two reactions that govern the Calvin–Benson cycle (R01063 and R01523) by sequestering the enzymes that catalyze these reactions, as discussed in (*25*). Additional details about individual AMGs are provided in Supplementary Text.

### Assessing AMG impacts using FBA

Here, following a parsimonious approach, we maximized the collective change in the flux in AMG reactions (taking care to enforce the correct direction of change) while simultaneously minimizing the flux through non-AMG reactions, subject to the constraint that the biomass flux remains near its optimal value. We considered a linear trade-off between the two objectives, parametrized by an *AMG reaction emphasis factor*, 𝛾. Higher values of 𝛾 favor maximizing AMG-promoted reaction flux changes, and lower values favor minimizing the non-AMG flux. We sorted each AMG-regulated reaction 𝑟 depending on the direct, unmediated impact of the AMG perturbations on its flux before (𝑣_𝑟_ ) and after (𝑣^′^ ) perturbation, as shown in Table 2. We also considered the set R_𝑔_ of all gene-regulated reactions (including those regulated by AMGs). After determining the optimal host flux distributions using the primary objective (Equations S1–S3), we then minimized the secondary objective 𝐹_𝑠𝑒𝑐_ (𝑣_𝑟_ ):

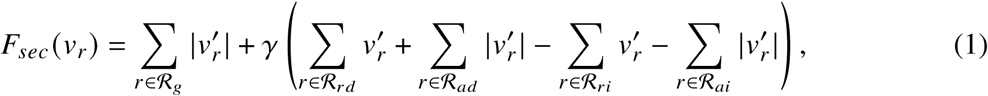

**Table 2:**
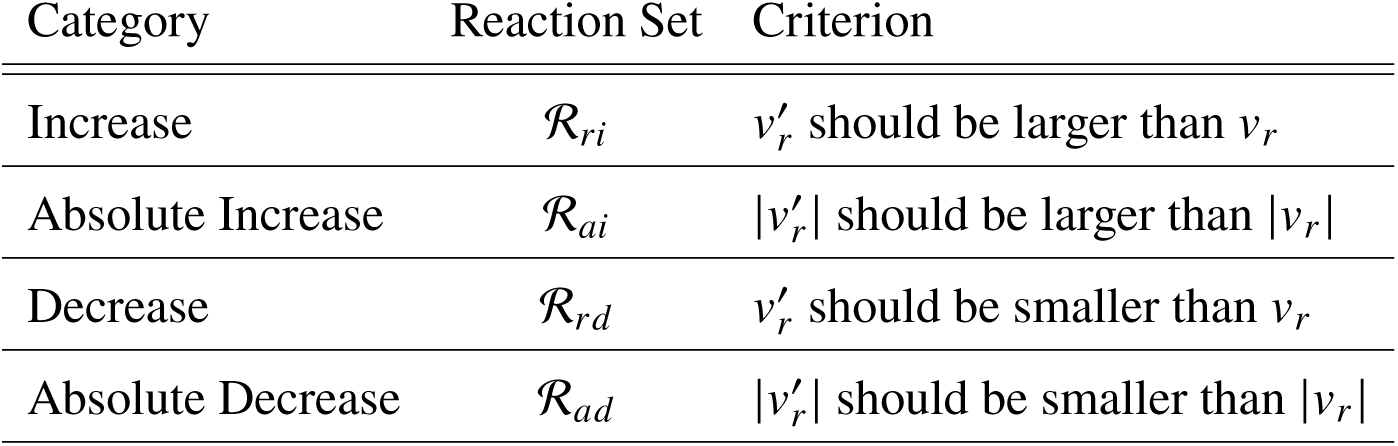
AMG-regulated reaction sets. Each reaction set is denoted R_𝑥𝑥_, where 𝑥𝑥 denotes the respective category reactions (𝑟𝑖, 𝑎𝑖, 𝑟𝑑, 𝑎𝑑), and can be categorized based on the change in flux through the reactions after infection perturbation.

subject to the constraints that i) the nutrient uptake does not increase too much (we use a factor of 2 as an upper limit, but also test with a factor of 5) and ii) the host biomass flux does not decrease below a specified threshold fraction (we use 50%) from the primary objective. We also retained any additional constraints used in determining the unperturbed flux distributions.

To evaluate how phage production limits host growth in various nutrient scenarios, we swept the full range of possible host growth rates for the given constraints on phosphate and ammonia uptake. We considered unconstrained uptake and uptake-limited scenarios where no more than 150%, 90%, or 20% of the uptake rates that yield optimal host growth can be achieved. Motivated by the reported impact of the AMG *phoH* (*12*), we also considered two scenarios in which the phosphate uptake is 3× or 30× greater than optimal for host growth. In each scenario, we considered 50 fixed values of host growth, evenly spaced between optimal growth in the scenario and near zero (0.0001%) growth. At each fixed value of host growth, we computed the maximum attainable phage replication rate, thereby constructing a Pareto front for the trade-off between host and phage growth. We repeated this process, restricting each AMG-hijacked reaction (except for phosphate exchange) to its target range to construct AMG-modulated trade-off curves.

### Subsystem mass flux graphs

We constructed subsystem mass flux graphs to elucidate the impacts of AMGs at a coarse-grained subsystem level. In these graphs, each node represents an annotated metabolic subsystem (e.g., biomass formation, photosynthesis, or oxidative phosphorylation). Reactions in the iSO595v7 model were annotated with a subsystem, though a plurality was not assigned a specific subsystem (here, we assigned these the *Unspecified* subsystem label). For a given reaction flux distribution, we computed the mass flux through a subsystem by first finding the net production or consumption rate of each metabolite by reactions in that subsystem. We accounted for metabolites that enter or exit the system (e.g., via exchange reactions or biomass secretion) by introducing an additional reaction subsystem, denoted ∅, corresponding to the environment that closes the system. Mass conservation requires that the mass-weighted sum of the positive metabolic production rates be equal in magnitude to the mass-weighted sum of the negative metabolic production rates (i.e., the consumption rates) in each reaction subsystem. We took this value to be the total flux through the subsystem. The mass flux between subsystems was calculated for each metabolite by determining the difference in mass production rates between two systems with directed edge weights being assigned the positive direction for mass flux.

To compare two subsystem mass flux graphs (e.g., for a healthy versus an infected cell), we constructed mass flux difference (MFD) and mass flux sum (MFS) graphs. These graphs are constructed with the difference or sum of weighted adjacency matrices for two subsystem mass flux graphs. Specifically, for two interventions, we computed 𝐺_𝑎𝑑𝑑_ as the MFS graph of the two individual MFD graphs relative to an intervention-free baseline and then computed the MFD graph 𝐺_𝑡𝑟𝑢𝑒_of the simultaneous interventions relative to the same baseline. The deviation from independent impacts is captured by the MFD of 𝐺_𝑡𝑟𝑢𝑒_ relative to 𝐺_𝑎𝑑𝑑_. As a simple summary of this graph, we computed its root mean square edge weight.

### Predicting the effects of CP12 on N-limited growth in MED4 and *S. elongatus*

In addition to the iSO595v7 MED4 metabolic model of (*37*) modified and considered throughout this work, we also analyze the iJB785 model (*50*), which we have modified by imposing a maximum RuBisCO utilization of 4.7 mmol gDW^−1^ h^−1^ following the method of (*37*); this does not affect the predicted growth rates in a standard medium but does prevent infinite growth in a replete medium. For both the RuBisCO-constrained iJB785 model and the iSO595v7 model, we conducted a parameter sweep of the N and P uptake constraints for the *cp12* mutant and WT. N and P uptake restrictions were achieved by constraining all the uptake of molecules containing either nutrient; note that MED4 lacks the ability to uptake nitrate. Based on Figure 1, we chose to set the flux reduction in the R01063 and R01523 reactions to 80%. We implemented low-light conditions as a 40% decrease in activity in photosystems I and II. We observed that the magnitude of the separation between high- and low-light conditions in Figure 7 is highly sensitive to the percentage reduction of R01063, but that this does not affect the qualitative behavior. In the results presented here, we selected a P uptake rate halfway through the parameter sweep (from 0 to maximum), which coincides with the value reported by (*50*). We computed the log2 ratio of the optimal growth rate for the mutant and WT simulations at each value of N-flux limitation and in the two light conditions considered. For comparison with experiment, we selected points near the transition from high to low log2 ratio (as a function of the N flux). We also calculated the Pearson correlation coefficients for these log2 ratios between experimental measurements and model simulations (Figure S9).

### Mutant engineering

We selected *Synechococcus elongatus* PCC 7942 as the expression chassis because it is a genetically tractable, model cyanobacterium with well-characterized neutral chromosomal integration sites and a growing toolkit of standardized integration (suicide) vectors and promoter systems (*62*). For heterologous gene insertion, we targeted Neutral Site II (NSII), employing a double homologous recombination (“suicide vector”) strategy with flanking NSII homology arms and a chloramphenicol resistance cassette, and placed the construct under the *Ptrc* promoter, which is broadly used in *S. elongatus* and other cyanobacteria for robust transcription (*63–65*). The *cp12* coding sequence was codon optimized for *S. elongatus* translational preferences to enhance expression, synthesized *de novo*, and cloned into the NSII suicide integration vector upstream of the chloramphenicol marker. The recombinant plasmid was transformed into *S. elongatus* PCC 7942, and chloramphenicol-resistant colonies were isolated. Complete segregation of the insertion was confirmed by genomic DNA extraction, PCR across the NSII locus, and Sanger sequencing to verify precise integration of the codon-optimized *cp12* cassette into the NSII site under the *Ptrc* promoter.

### Experimental validation of the CP12 effect on *S. elongatus* growth under N-limited conditions

Both WT and *cp12* engineered *S. elongatus* PCC 7942 strains were maintained in 1.2 L Roux culture flasks containing 0.4–0.6 L of BG-11 medium at 24 ± 2 ^◦^C. Cultures were illuminated with a photosynthetic photon flux density (PPFD) of approximately 100 µmol m^−2^ s^−1^ using Monios-L LED Full Spectrum Grow Lights, measured with an LI-250A Light Meter equipped with an LI-190SA Quantum Sensor (LI-COR, Inc.). Mixing was achieved using polytetrafluoroethylene (PTFE) octagon spinbar magnetic stirring bars (9.5 × 25.4 mm) at approximately 200 rpm. Cultures were continuously sparged with nitrogen gas containing 2% CO_2_ and were diluted to an optical density at 750 nm (OD_750_) of 0.2 every 3–4 days to maintain the exponential growth phase. Twenty-four hours prior to the nitrogen limitation experiment, *S. elongatus* cells were transferred to a nitrogen-free BG-11 medium (BG-11/-N) by centrifugation at 6, 000 × 𝑔 for 5 min, followed by resuspension in BG-11/-N to remove residual nitrogen. Growth assays were performed in 96-well flat-bottom microtiter plates with a 200 µL culture volume and four biological replicates per treatment. The target starting OD_750_ was 0.05 for all growth assays. Nitrogen was supplied as NaNO_3_ at concentrations of 1500, 1200, 800, 400, 200, 120, 60, 30, 15, and 0 mg/L. Cultures were grown under two PPFDs: 50 µmol m^−2^ s^−1^ and 230 µmol m^−2^ s^−1^ in a Labtron Light incubator with a 2% CO_2_ atmosphere. The cell density was monitored by measuring OD_750_ using a BioTek Synergy Neo2 Hybrid Multimode Microplate Reader (Agilent Technologies). Growth measurements were taken at regular intervals over the experimental period to generate growth curves and calculate growth parameters.

### Growth curve data analysis

To analyze the growth curve data (see Figure S8), we first subtracted the OD_750_ values of the blank wells from each data point. We then fit a logistic curve to each replicate OD_750_ time series:

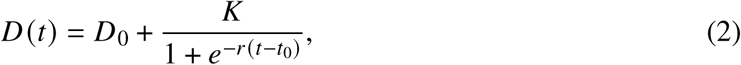

where 𝐷(𝑡) is optical density as a function of time, 𝐷_0_ is the baseline optical density, 𝑟 is the logistic growth rate, and 𝐾 is the logistic carrying capacity. We exclude points that occur after the maximum value is reached to avoid fitting to population collapse after nutrient depletion. We perform the fit using parameter bounds informed by the maximum and minimum observed values and applying the trust region reflective algorithm implemented in the Python library SciPy, which also provides error estimates of each parameter. After fitting each growth curve, it is straightforward to compute a maximum growth rate by setting 𝑃(𝑡) = 𝐷(𝑡) − 𝐷_0_ and using the fact that 𝐷^·^ to find that 𝐷^·^ is maximized at a value of

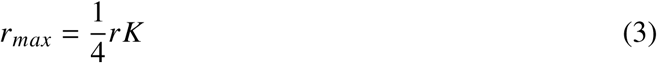

when 𝑃 = <u>^1^</u> 𝐾. We propagate the errors from the fitting procedure using a linearization approximation. We discard replicates with growth rate estimates whose error exceeds half the value of the estimate; this typically occurs when the change in the optical density is very small. To compute the difference in log2 maximum growth rates, we consider replicates in pairs based on their distance from the edge of the well plates to better control for carbon limitation and evaporation.

## Footnotes

1 Because a reaction can be mediated by several genes and a gene can mediate several reactions, the number of essential reactions need not be equal to the number of essential genes.

2 PSIIabs represents the photon absorption in photosystem II.

3 FAKEOrthophosphateEX is the phosphate exchange reaction; the FAKE prefix indicates that it was added through gap filling by (*38*).

## Acknowledgments

We acknowledge the insightful discussions with Martin H. W. Gruebele, Zaida Ann Luthey-Schulten, and Aaron Chan from the University of Illinois Urbana-Champaign; John Casey from Lawrence Livermore National Laboratory; and William Nelson, Jason McDermott, and Chris Oehmen from Pacific Northwest National Laboratory.

## Author contributions

**Conceptualization:** SF, RW; **Methodology:** SF, JCR, WS; **Software:** SF, JCR, WS; **Formal Analysis:** JCR, WS, SF; **Investigation:** SF, JR, WS, PB; **Resources:** SF, WS, DK, JQ; **Data Curation:** WS, SF, JR; **Writing - Original Draft Preparation:** JCR, WS, SF, PB, APM; **Writing - Review & Editing:** All authors; **Visualization:** JR, WS, SF; **Project Administration:** SF, WQ; **Funding Acquisition:** MC, DP, WQ.

## Supplementary materials

Supplementary Text Figures S1 through S5 Tables S1 through S3 Software S1 (available at https://github.com/NWBRaVE/p-hm2-amgs-in-med4). Data S1

## Supplementary Materials for

### Other Supplementary Materials for this manuscript

Software S1 (also available at https://github.com/NWBRaVE/p-hm2-amgs-in-med4). Data S1

### Supplementary Text

Additional details about AMGs in *P. marinus*

In *Prochloroccocus marinus*, CP12 inhibits the Calvin–Benson cycle light-limited conditions to shift the metabolism in favor of the pentose phosphate pathway (PPP) for nucleotide synthesis (*25*). Also favoring the PPP, the *cobS* auxiliary metabolic gene (AMG) affects the reaction responsible for cobalamin (B12) synthesis. Cobalamin is predicted to increase ribonucleotide diphosphatase function in the nucleotide metabolism, thus increasing DNA synthesis (*43*). Similarly working in favor of nucleotide synthesis, the AMG *talC* codes for transaldolase, an enzyme active in the PPP (*25*).

The next set of AMGs has the largest correlation with the photosynthesis pathway in *Prochloroccocus marinus*. The *psbA* and *psbD* AMGs correlate with the D1 and D2 protein subunits, respectively, at the photosystem II reaction center. The D1 and D2 subunits bind the primary electron transfer chain along with the P680 reaction center chlorophyll special pair (*48*). These AMGs are predicted to have a large effect on energy binding in photosynthesis. The *ho1* AMG catalyzes the reaction responsible for the production of biliverdin, a precursor to light-harvesting pigments (*47*). Similarly, the *cpeT* and *cpcT* AMGs catalyze the production of the light-harvesting pigments phycocyanobilin (*44*). Further, the *pebS* AMG codes for an enzyme that produces the light-harvesting pigment phycoerythrobilin (*44*). These AMGs work to increase the pigments needed for photosynthesis. Along with the AMGs predicted to increase photosynthesis, the *hlip* and *hsp20* AMGs code for proteins that dissipate heat at the photosynthesis reaction center (*45*). These last two genes do not directly participate in the host metabolism; however, these AMGs may biologically play a thermodynamic role.

The function of *mazG* as an AMG is relatively unknown. In other organisms, it is predicted to regulate levels of (p)ppGpp, a putative alarmone, and trick the organism into a nutrient-replete state. This would cause the host to use less nutrients and allow the phage to increase replication (*28*). In (*66*), it is argued that *mazG* may degrade nucleotides to decrease GC content. During the infection of *S. elongatus*, this may favor the phage—*S. elongatus* has a GC content near 60%, while cyanophages have GC content close to 40% (P-HM2 has a GC content of 38%). In contrast, MED4 has an even lower GC content of just over 31%, so the role of *mazG* in this host is less clear. Additionally, the function of the *phoH* AMG remains relatively unclear. There is evidence that it functions as a phosphate uptake gene and codes for ATPase enzymes (*12*). However, *phoH* is significantly upregulated during phage infection, and this could cause an increase in phosphate available for phage replication (*12*).

#### Flux balance analysis (FBA)

Flux analysis can be used to extract testable, mechanistic predictions from limited data (*30*). Reaction rate parameters are often unknown or poorly constrained, making a kinetic model nonviable. In contrast, the stoichiometric relationships encoded by chemical reaction networks are often tightly constrained.

For a set of metabolites 𝑀 governed by reactions 𝑅, flux balance analysis (FBA) exploits stoichiometric constraints to determine the steady-state flux distributions that maximize a biological objective 𝐹_𝑜𝑏𝑖_ (𝑣_𝑟_ ). The *reaction flux* is the instantaneous rate of a reaction 𝑟 ∈ 𝑅 per unit cell mass, here denoted 𝑣_𝑟_ ∈ R (typically measured in units of millimoles per gram of dry weight per hour). The rate per unit cell mass at which a metabolite 𝑚 ∈ 𝑀 is produced (or consumed) by a reaction 𝑟 ∈ 𝑅 is given by the stoichiometric coefficient, 𝑆_𝑟,𝑚_, times the flux 𝑣_𝑟_ of 𝑟.

FBA is formulated as a linear optimization problem:

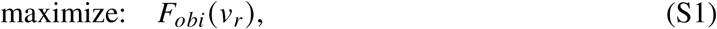

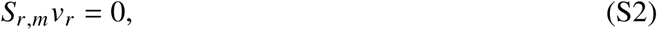

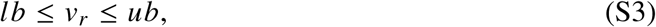

where a biologically inspired objective function, 𝐹_𝑜𝑏𝑖_ (𝑣_𝑟_ ), such as a biomass reaction encoding the metabolic components necessary to produce one unit (typically one gram) of biomass, is maximized to represent cell growth. The steady-state assumption enforces the 𝑆_𝑟,𝑚_𝑣_𝑟_ = 0 constraints on the flux values, and the reaction flux bounds, lower bounds 𝑙𝑏 ∈ R_≥0_, and upper bounds 𝑢𝑏 ∈ R_≥0_ can be used to constrain the solution. Solutions to 𝑆_𝑟,𝑚_𝑣_𝑟_ = 0 that satisfy all specified constraints are called *feasible*; those that do not are called *infeasible*.

Secondary objectives can supplement FBA for more complex models. In *parsimonious* FBA (pFBA), the secondary objective is to minimize the total (absolute) flux of enzyme-catalyzed reactions. Thus, pFBA produces solutions that maximize the biomass while simultaneously minimizing the enzyme production. In *flux variability analysis* (FVA), each reaction flux is taken, in turn, to be an objective to minimize or maximize, providing the minimum and maximum fluxes that each reaction can support while maintaining optimal growth (with a chosen tolerance). In this study, we have performed pFBA and FVA using COBRApy (*42*) with the GNU Linear Programming Kit (GLPK) back end (*67*).

Viral infection can be modeled as a perturbation to the host system. It has been demonstrated experimentally that gene knockout perturbation response often minimizes the change in flux rather than maximizes the biomass production (*29, 32*). When considering the impact of AMG-hijacked reactions individually or pair-wise, we specify a target range for each hijacked reaction by taking 10% of the interval in which phage production can be supported above half its maximum value (see Figure 1). Following (*29*), to compare to a baseline flux distribution, we restrict the AMG-hijacked reaction(s) to the corresponding target range(s) and perform minimization of metabolic adjustment (MOMA) to find the flux distribution that satisfies the new constraints and is closest to the baseline (by the L1 norm).

#### Structural sensitivity analysis for model comparison

In metabolic modeling, structural sensitivity analysis is used to show the network response to a flux perturbation in one or more reactions. Structural sensitivity analysis uses a matrix of reaction stoichiometric coefficients, 𝑆, and a matrix of reference flux values, 𝑣 (*68*). We used FBA to generate 𝑣 from the assumed steady state 𝑆𝑣 = 0. To analyze how the network responds to a perturbation 𝛿 in reaction 𝑘, we can perturb the flux as 𝑣_𝑘_ + 𝛿_𝑘_ (*68*). We find the minimal adjustments to satisfy

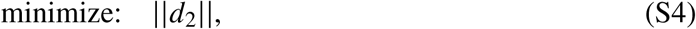

subject to:

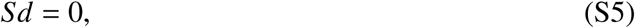

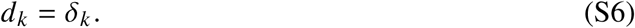

In this case, the sensitivity, 𝑥, of reaction 𝑖 relative to the perturbation in reaction 𝑘 is

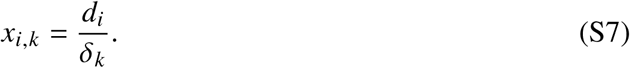

This method can be extended to compare models by applying the perturbations, 𝛿, to both models. This gives the network sensitivities of both organism models to the same perturbation. Using a subset of reactions that are found in both models, we can find the Pearson correlation for sensitivities to measure the similarity of the networks’ responses to the perturbation (*69*).

#### AMG cyanophage cloning process

Cyanophage protein coding sequences (CDSs) were codon optimized for *S. elongatus* expression using Benchling and ordered as IDT gBlocks. Cyanophage protein CDSs were inserted into pCV0055 via Gibson assembly. pCV (*70*) was linearized using oligonucleotides JQ0045/JQ0061, and cyanophage protein CDSs were amplified using oligonucleotides JQ0057/0058. Cyanobacteria were transformed as previously described by the Golden lab (*71*). Briefly, 15 mL of exponential phase WT Syn7942 per transformation are collected and pelleted at 3700 rpm for 5 min. The pellet is washed once with room-temperature sterile 10 mm NaCl and resuspended in 150 µL of BG11 media per transformation in sterile 1.5 mL Eppendorf tubes. Then, 150 ng of plasmid is added to each transformation and left to incubate at 30 ^◦^C overnight protected from light. The following morning, all 150 µL are plated onto BG11 + 1.5% agar + 5 µg/mL chloramphenicol. After colonies appear (∼4 days), the clonal integration zygosity is determined by colony PCR using oligonucleotides JQ0081/JQ0082. Clones that are homozygous for integration and return the correct payload integration by Sanger sequencing are expanded and frozen for long-term storage as previously described (*72*).

---

**Figure S1:**
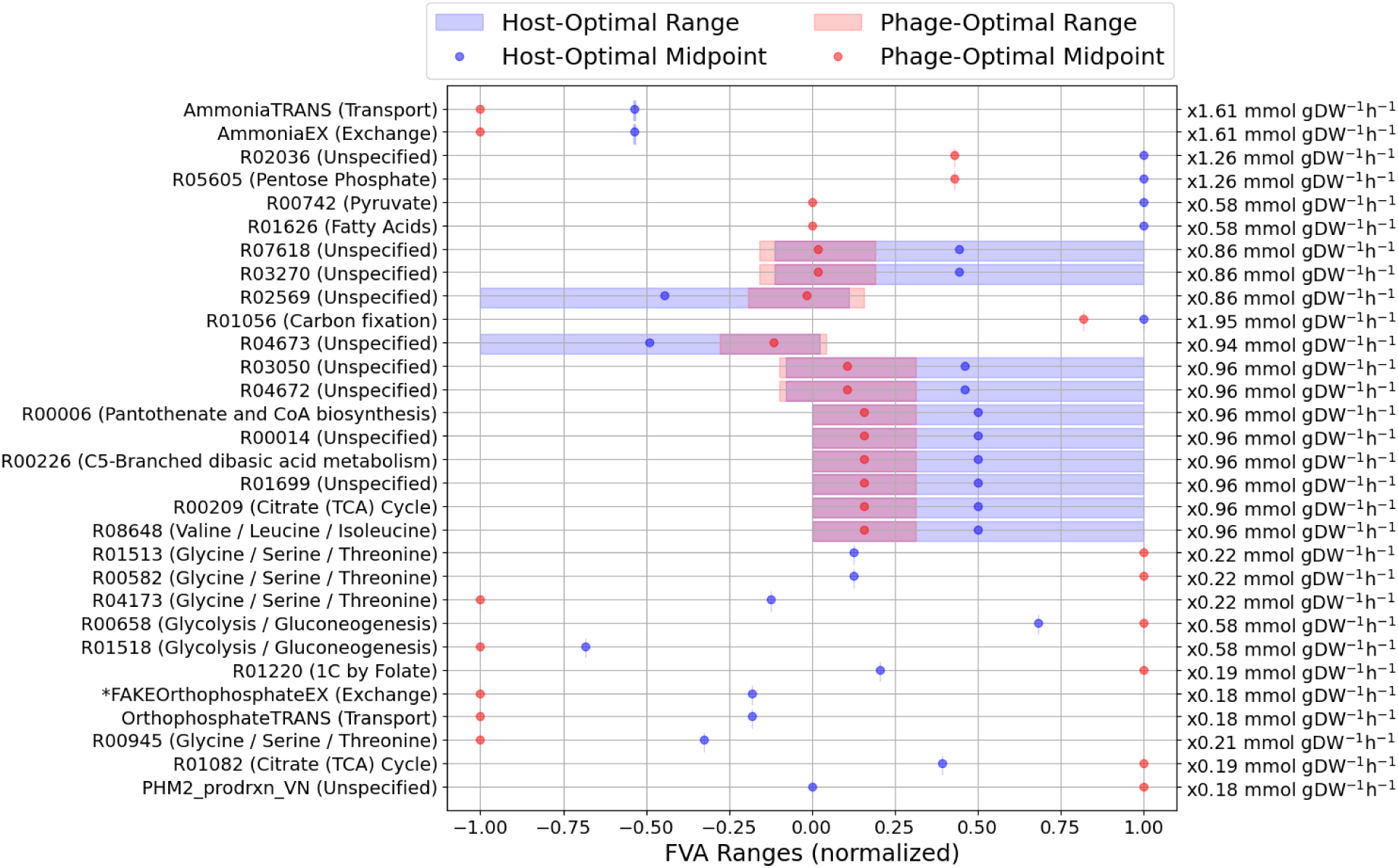
FVA ranges for 30 reactions most affected by optimizing for phage versus host biomass. Blue ranges indicate the flux variability when maximizing for the host biomass, while red ranges are for the variability associated with maximizing the phage biomass. Any overlap is shown in purple. Ranges are normalized by the most extreme feasible flux across the two scenarios (normalization factors depicted along the right-hand side of the figure). Reactions with normalization below that of phage production (PHM2 prodrxn VN) are not included in this analysis. Reaction IDs and subsystems are given along the left-hand side. Reactions are ordered by the (unnormalized) midpoint distance. AMG-hijacked reactions are indicated with an asterisk.

**Figure S2:**
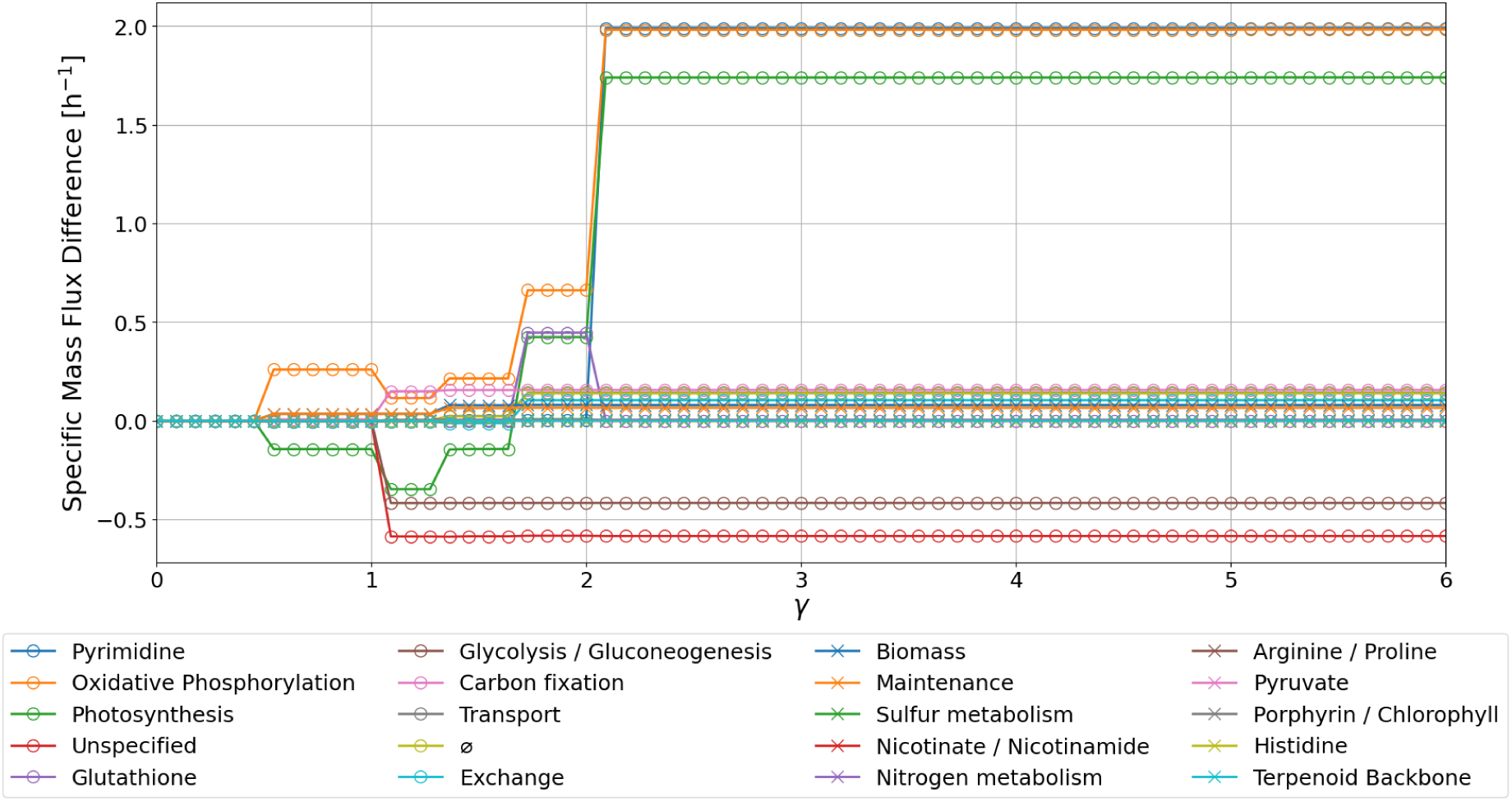
Changes in the specific mass flux through the 20 most impacted subsystems. Results are shown as a function of the AMG reaction emphasis factor, 𝛾. The ten subsystems with the largest responses are marked with circles, the remaining ten use an X symbol. Increasing 𝛾 more heavily weights changing the flux of AMG-hijacked reactions relative to minimizing the flux through other gene-regulated reactions (see Methods). In contrast to Figure 4, the vertical axis uses a linear scale.

**Figure S3:**
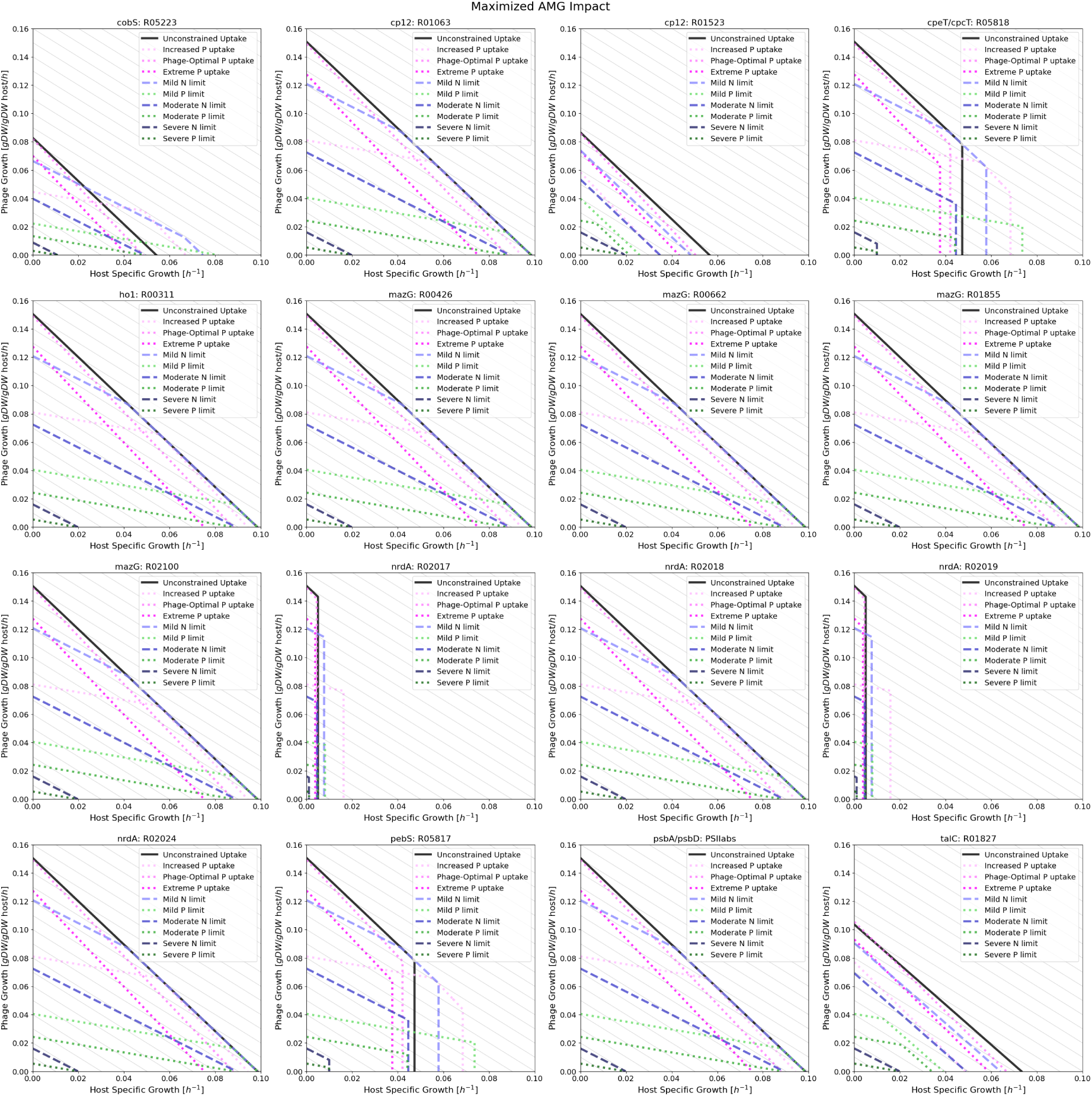
Phage–host biomass Pareto fronts. Full-size image available online. Each panel depicts the instantaneous relationships between the maximum (exponential) host-specific growth rate and the maximum (linear) phage growth rate for an AMG-hijacked reaction, as indicated in the figure titles. We show the Pareto fronts for several nutrient-limiting scenarios: unconstrained uptake and mild/moderate/severe limiting of N or P uptake. Mild, moderate, and severe limits correspond to 150%, 90%, and 20% of the values required for maximal host biomass production, respectively. When growth is not constrained by the nutrient uptake, it is instead limited by the RuBisCO efficiency. In addition, we also considered two scenarios in which the cell is forced to uptake more phosphate than is optimal (Increased P uptake: 3× host-optimal uptake; Phage-optimal P uptake: 6× host-optimal uptake; Extreme P up_S_ta_8_ke: 12× host-optimal uptake).

**Figure S4:**
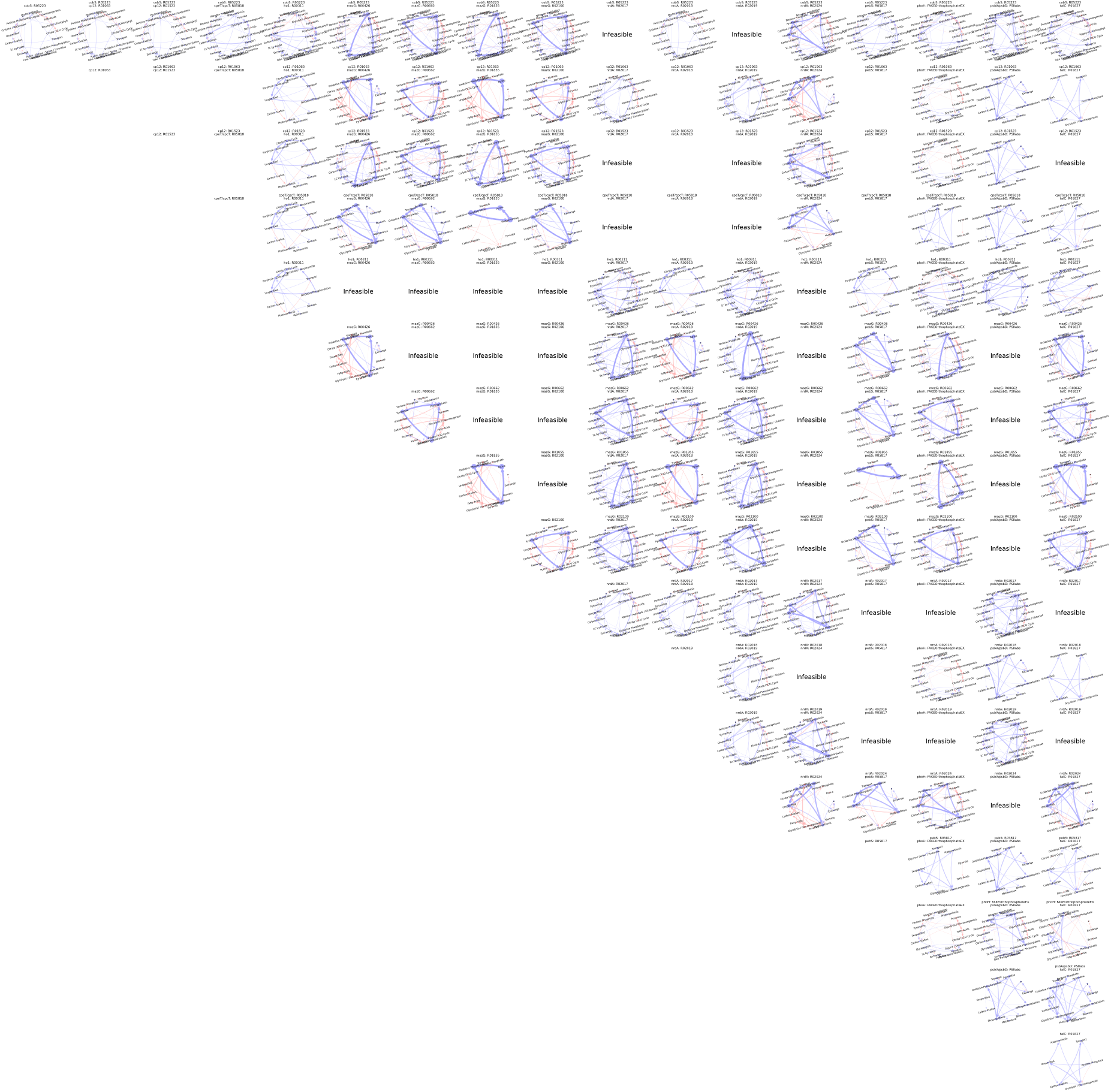
Pairwise AMG-hijacked reaction impacts on subsystem fluxes. Full-size image available online. Each column and row correspond to an AMG-hijacked reaction. Along the diagonal, a single reaction is hijacked. Along the upper off-diagonals, pairs of reactions are hijacked. The resulting mass flux difference graphs (see Methods) are plotted, with blue (red) indicating increased (decreased) mass flux (changes below 0.01h^−1^ are not shown). The reactions hijacked are indicated with text labels on each graph plot. When simultaneous hijacking is not possible because of stoichiometric constraints, the text “Infeasible” appears in place of a graph plot.

**Figure S5:**
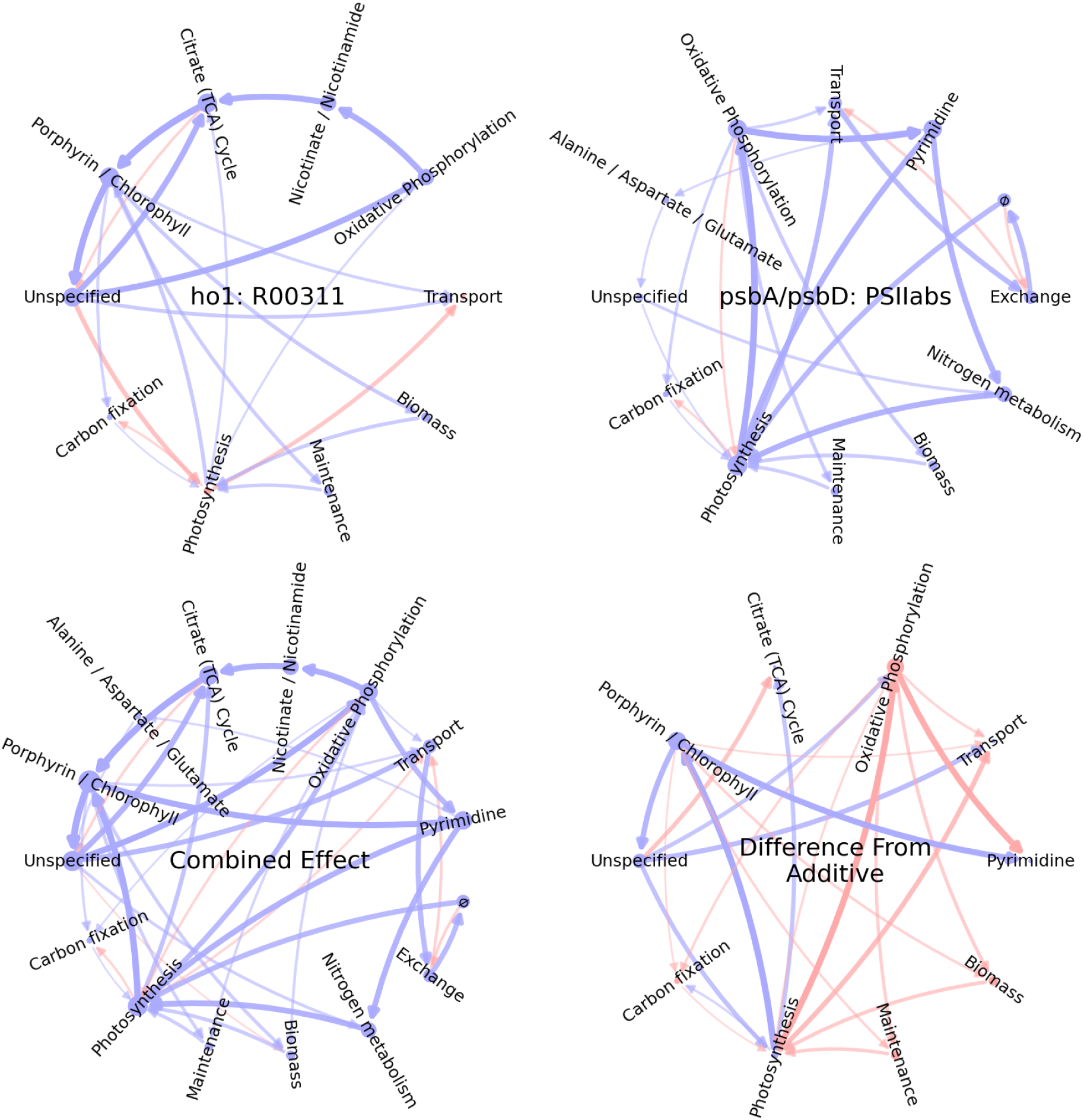
Deviation from the additive impact of two AMG-hijacked reactions. In the top row, the mass flux difference (MFD) graphs for PSIIabs and R00311 are shown. The bottom left graph is the MFD graph (see Methods) for their combined impact. The bottom right graph shows the deviation of the combined impact MFD from a purely additive impact. For example, the red edge from Oxidative Phosphorylation to Photosynthesis denotes that hijacking both reactions has a smaller impact on this flux than the sum of the individual impacts. In all graphs, edges with specific mass flux changes below 0.01 h^−1^ are not shown, nor are nodes that lack incoming or outgoing edges above this threshold.

**Figure S6:**
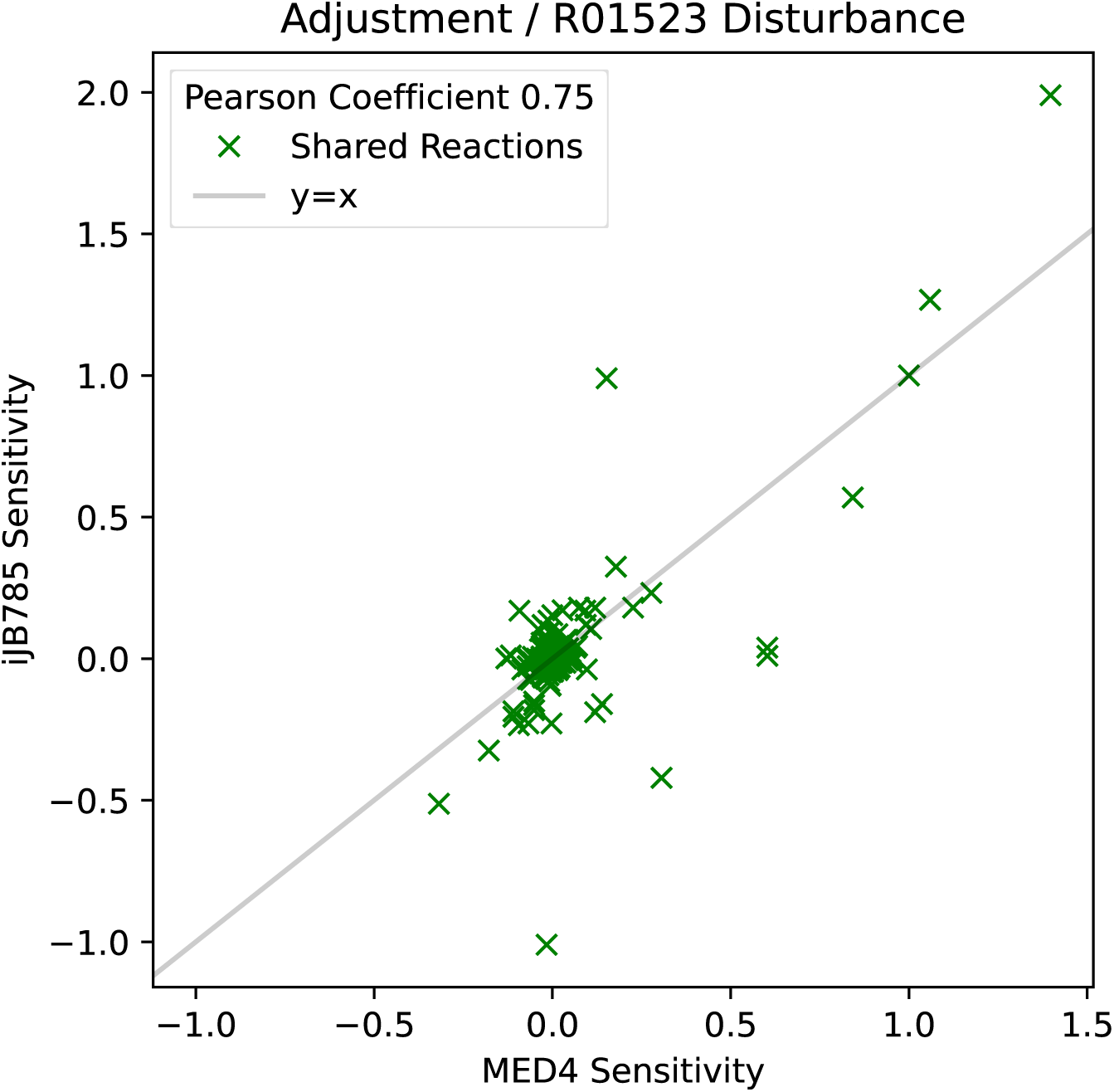
Sensitivity analysis of all shared reactions relative to the disturbance of CP12-hijacked reaction R01523 in both iJB785 and iSO595v7. The green “×” symbols indicate shared reactions between the two models. The gray line shows the perfect correlation line. The sensitivities of all the shared reactions show a high correlation, with Pearson coefficient of 0.75.

**Figure S7:**
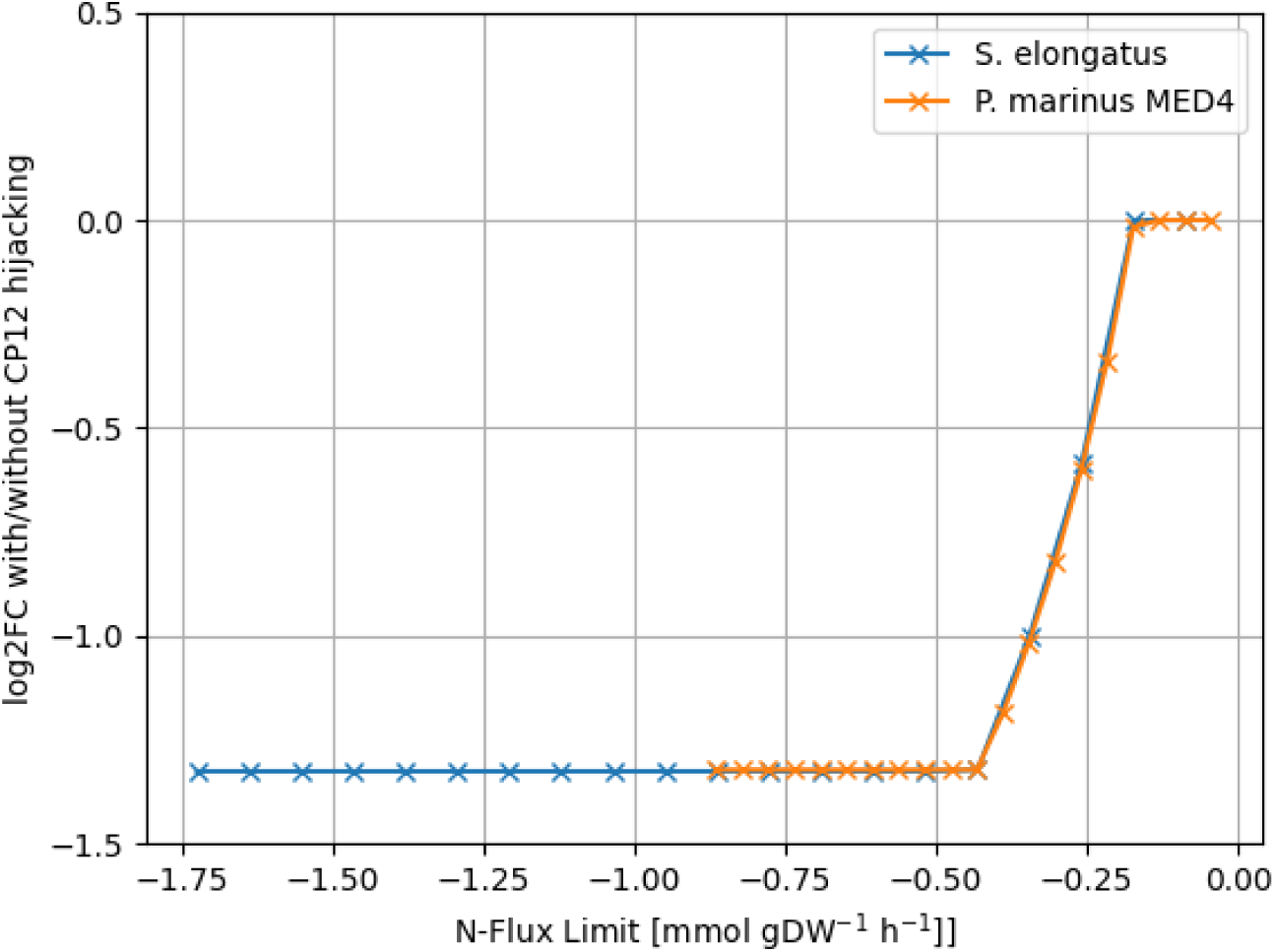
Impact of CP12 on N-limited growth in iJB785 and iSO595v7. The log2 optimal growth ratio between the CP12-hijacked and unmodified FBA models is plotted for various values of the maximum N flux (more negative means more N uptake). The values for the iJB785 model of *S. elongatus* are shown in blue, and the values for the iSO595v7 model of MED4 metabolism are shown in orange. Phosphate uptake was constrained to half the replete flux value. Photosystem activity was not constrained. The effect of *cp12* is modeled as a reduction of 80% in the flux through reactions R01063 and R01528. To ensure comparability between the models, the larger of the minimum unconstrained fluxes in the two models was used to constrain both models. The difference in the horizontal scale arises from the ability of *S. elongatus* to acquire N as both nitrate and ammonium, whereas MED4 cannot uptake nitrate. Nevertheless, the differential response to the N limitation in the presence of constitutively active *cp12* is nearly identical in the two models.

**Figure S8:**
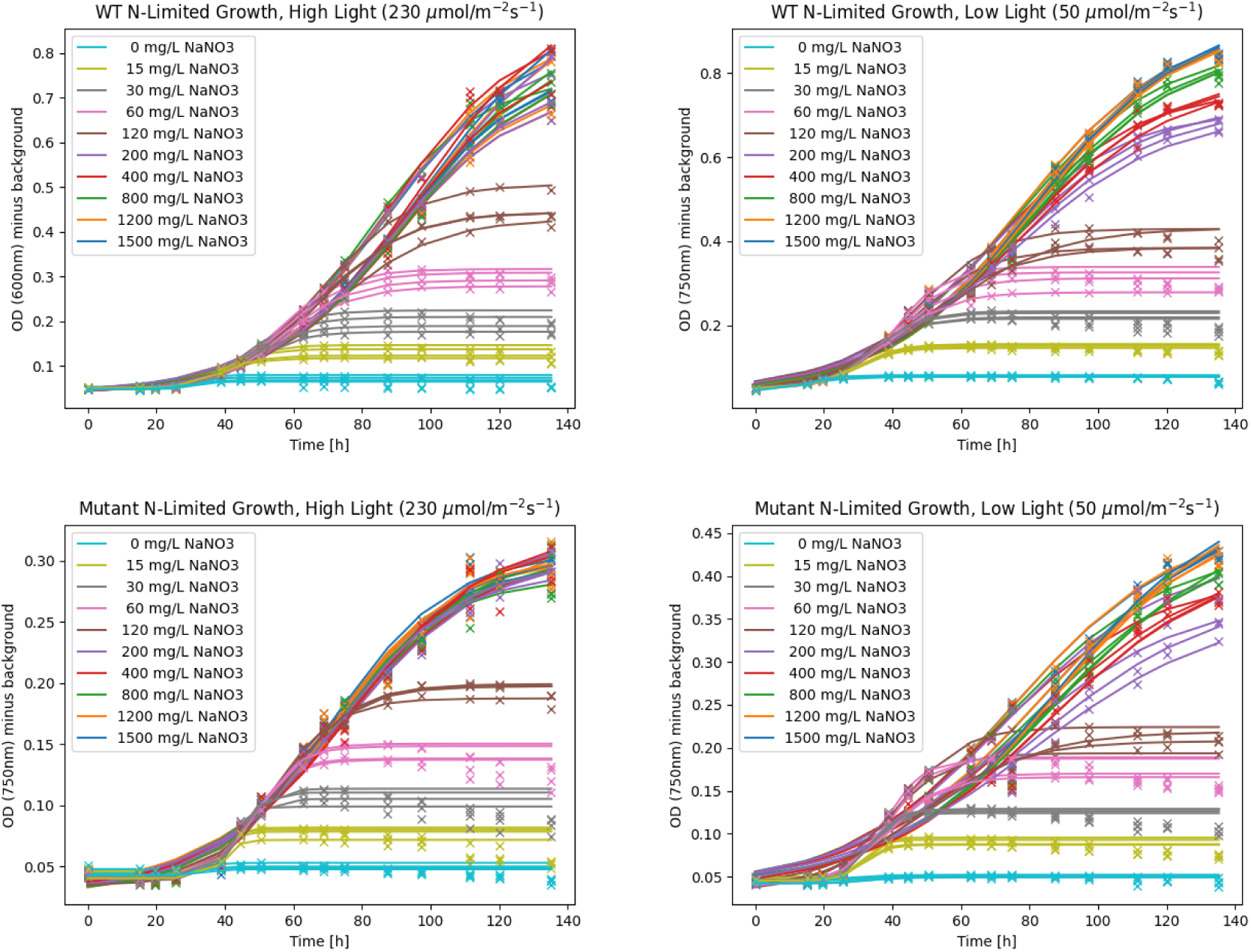
Growth curves from N-limited growth experiments. Data points are indicated by crosses, and logistic fits to each replicate are indicated by solid lines. Curve fits disregard the portion of the trajectory during which OD_750_ is decreasing. Line and cross colors indicate the concentration of NaNO_3_ added to the growth medium during preparation, which serves to constrain N availability. In the top row, OD_750_ growth curves are presented for WT *S. elongatus* in high and low light, while the bottom row depicts growth curves for the mutant *S. elongatus* with constitutively active *cp12* in the same two light conditions.

**Figure S9:**
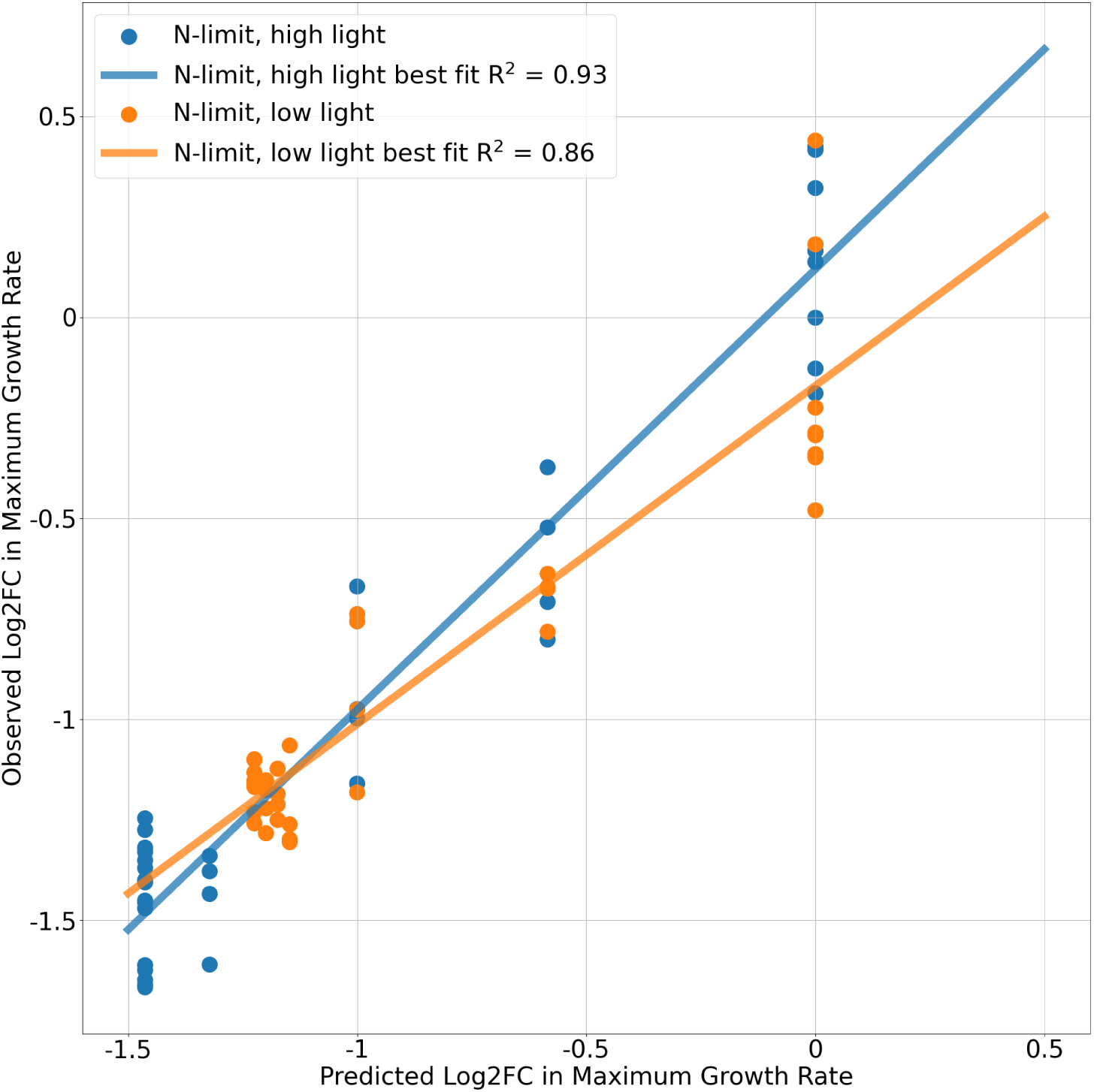
Correlation between experimentally measured growth rate changes and simulated growth rate changes from the model. The log2 optimal growth ratio between the CP12-hijacked and unmodified FBA models is plotted for both the experimental measurements (y-axis) and model simulations (x-axis). The values for the ”N-limit, high light” conditions are shown in blue, and the values for the ”N-limit, low light” conditions are shown in orange. Pearson correlation coefficients are calculated for both high light (0.93) and low light (0.86) conditions.

**Table S1:** Chemical reaction expressions and subsystems for AMG-hijacked reactions.

| Gene: Reaction ID | Reaction Expression | Subsystem |
| --- | --- | --- |
| cobS: R05223 | [Adenosine-GDP-cobinamide] + [alpha-Ribazole] →<br>[Cobamide coenzyme] + [GMP] + [H<sup>+</sup>] | Porphyrin / Chlorophyll |
| cp12: R01063 | [D-Glyceraldehyde 3-phosphate] + [NADP<sup>+</sup>] + [Orthophosphate] ⇌<br>[3-Phospho-D-glyceroyl phosphate] + [H<sup>+</sup>] + [NADPH] | Unspecified |
| cp12: R01523 | [ATP] + [D-Ribulose 5-phosphate] ⇌<br>[ADP] + [D-Ribulose 1,5-bisphosphate] | Carbon fixation |
| cpeT/cpcT: R05818 | [Biliverdin] + [Reduced ferredoxin] →<br>[15,16-Dihydrobiliverdin] + [Oxidized ferredoxin] | Porphyrin / Chlorophyll |
| ho1: R00311 | [Heme] + 3.0 [Oxygen] + 3.0 [Plastoquinol] →<br>[Biliverdin] + [CO] + [Fe<sup>2+</sup>] + 3.0 [H<sub>2</sub>O] + 3.0 [Plastoquinone] | Porphyrin / Chlorophyll |
| mazG: R00426 | [GTP] + [H<sub>2</sub>O] →<br>[Diphosphate] + [GMP] | Purine |
| mazG: R00662 | [H<sub>2</sub>O] + [UTP] →<br>[Diphosphate] + [UMP] | Pyrimidine |
| mazG: R01855 | [H<sub>2</sub>O] + [dGTP] →<br>[Diphosphate] + [dGMP] | Purine |
| mazG: R02100 | [H<sub>2</sub>O] + [dUTP] →<br>[Diphosphate] + [dUMP] | Pyrimidine |
| nrdA: R02017 | [ADP] + [Thioredoxin] →<br>[H<sub>2</sub>O] + [Thioredoxin disulfide] + d[ADP] | Purine |
| nrdA: R02018 | [H<sub>2</sub>O] + [Thioredoxin disulfide] + [dUDP] ⇌<br>[Thioredoxin] + [UDP] | Unspecified |
| nrdA: R02019 | [GDP] + [Thioredoxin] →<br>[H<sub>2</sub>O] + [Thioredoxin disulfide] + d[GDP] | Purine |
| nrdA: R02024 | [CDP] + [Thioredoxin] →<br>[H<sub>2</sub>O] + [Thioredoxin disulfide] + d[CDP] | Pyrimidine |
| pebS: R05817 | [Biliverdin] + [Reduced ferredoxin] →<br>[(3Z)-Phycocyanobilin] + [Oxidized ferredoxin] | Porphyrin / Chlorophyll |
| phoH: FAKEOrthophosphateEX | [Orthophosphate] ⇌<br>∅ | Exchange |
| psbA/psbD: PSIIabs | [Photon] →<br>[Photon absorbed by PSII] | Photosynthesis |
| talC: R01827 | [D-Glyceraldehyde 3-phosphate] + [Sedoheptulose 7-phosphate] ⇌<br>[D-Erythrose 4-phosphate] + [D-Fructose 6-phosphate] | Pentose Phosphate |

**Table S2:** Biomass production reaction for P-HM2. The phage mass is modeled as the mass deficit in this reaction due to amino acid and dNTP consumption. Stoichiometric coefficients and reaction velocity units are such that a reaction rate of 1 corresponds to 1 gram of phage produced per hour per gram dry weight of host.

| Amino Acid Consumed | mmol/g virus | Nucleotide Consumed | mmol/g virus |
| --- | --- | --- | --- |
| Alanine | 00.428 | ATP | 20.455 |
| Arginine | 00.195 | CTP | 00.005 |
| Asparagine | 00.312 | GTP | 00.006 |
| Aspartate | 00.300 | UTP | 00.007 |
| Cysteine | 00.023 | dATP | 00.378 |
| Glutamate | 00.247 | dCTP | 00.233 |
| Glutamine | 00.180 | dGTP | 00.233 |
| Glycine | 00.422 | dTTP | 00.378 |
| Histidine | 00.050 | Nucleotide Produced | mmol/g virus |
| Isoleucine | 00.269 | AMP | 00.009 |
| Leucine | 00.310 | ADP | 20.446 |
| Lysine | 00.192 | CMP | 00.005 |
| Methionine | 00.090 | GMP | 00.006 |
| Phenylalanine | 00.185 | UMP | 00.007 |
| Proline | 00.164 | Small Molecule Consumed | mmol/g virus |
| Serine | 00.389 | H<sub>2</sub>O | 20.446 |
| Threonine | 00.451 | Small Molecule Produced | mmol/g virus |
| Tryptophan | 00.039 | H | 20.446 |
| Tyrosine | 00.196 | Orthophosphate | 20.446 |
| Valine | 00.321 | Diphosphate | 01.249 |

**Table S3:** ***Prochloroccocus marinus* and P-HM2 biomass components.** The RNA mass for P-HM2 corresponds to the mRNA synthesis needed for the transcription of genes encoding structural proteins and does not directly contribute to the phage mass in this model.

| Component | MED4 (g/gDW host) | P-HM2 (g/gDW virus) |
| --- | --- | --- |
| Protein | 0.6492 | 0.6007 |
| DNA | 0.0131 | 0.3993 |
| RNA | 0.0525 | 0.0132 |
| Lipids | 0.1284 |  |
| Other | 0.1824 |  |

